# Cryptic genetic variation defines the adaptive evolutionary potential of enzymes

**DOI:** 10.1101/232793

**Authors:** Florian Baier, Nansook Hong, Gloria Yang, Anna Pabis, Alexandre Barrozo, Paul D Carr, Shina CL Kamerlin, Colin J Jackson, Nobuhiko Tokuriki

**Author notes:** Corresponding author: Nobuhiko Tokuriki, Michael Smith Laboratories, University of British Columbia, Vancouver, V6T 1Z4, BC, Canada.

## Abstract

Genetic variation among orthologous proteins can cause cryptic phenotypic properties that only manifest in changing environments. Such variation may also impact the evolutionary potential of proteins, but the molecular basis for this remains unclear. Here we perform comparative directed evolution in which four orthologous metallo-β-lactamases were evolved toward a new function. We found that genetic variation between these enzymes resulted in distinct evolutionary outcomes. The ortholog with the lower initial activity reached a 20-fold higher fitness plateau exclusively *via* increasing catalytic activity. By contrast, the ortholog with the highest initial activity evolved to a less-optimal and phenotypically distinct outcome through changes in expression, oligomerization and activity. We show that the cryptic molecular properties and conformational variation of residues in the initial genotypes cause epistasis, thereby constraining evolutionary outcomes. Our work highlights that understanding the molecular details relating genetic variation to protein functions is essential to predicting the evolution of proteins.

## Introduction

Genetic diversity across orthologous proteins is thought to be predominantly neutral with respect to their native, physiological function, but can cause “cryptic genetic variation”, *i.e*., variation in other non-physiological phenotypic properties^1–3^. Cryptic genetic variation has been shown to play an important role in evolution, because genetically diverse populations are more likely contain genotypes with a “pre-adapted” phenotype, *e.g*., a latent promiscuous function, that confer an immediate selective advantage when the environment changes and a new selection pressure emerges^4–8^. Beyond these examples, however, we have little understanding of how genetic variation affects long-term adaptive evolutionary potential or the “evolvability” of proteins^9^. Many biological traits, in particular promiscuous proteins functions, often evolve by accumulating multiple adaptive mutations before reaching a fitness plateau or peak. Thus, the degree to which the trait can improve and the level of peak fitness that it can reach ultimately determines the evolutionary potential and outcomes^10^. Recent studies have shown that, due to intramolecular epistasis^11–13^ the phenotypic effects of the same mutations introduced into orthologous proteins can exhibit variable phenotypic effects^14,15^, this is true even if they are introduced into a genotype that differs by only a few other mutations^16,17^. However, having examined only a small subset of mutations, the degree to which genetic starting points determines long-term evolutionary outcomes given the same selection remains unclear^18,19^. Moreover, we have very little understanding of the molecular mechanisms that underlie the relationship between genetic variation and evolvability. It has been suggested that protein fold^20–22^ and protein stability^23–25^ can determine evolvability, but these alone cannot explain the prevalence of epistasis and the enormous variation we observe in the evolvability of different genotypes.

Metallo-β-lactamases (MBLs) encompass a genetically diverse enzyme family that confer β-lactam antibiotic resistance to bacteria (**Fig. 1a,b**)^26,27^. Our previous work demonstrated that most MBLs exhibit phosphonate monoester hydrolase (PMH) activity (*k*_cat_/*K*_M_) in the range of 0.1 to 10 M^−1^s^−127,28^. β-lactamase and PMH hydrolysis differs by scissile bond (C-N *vs*. P-O) and transition state geometry (tetrahedral *vs*. trigonal bipyramidal), making the PMH activity a catalytically distinct promiscuous activity for these enzymes. Here, we perform an empirical test of enzyme evolvability by performing directed evolution with orthologous MBLs (NDM1, VIM2, VIM7, and EBL1) towards PMH activity. Through detailed analysis of the genotypic and phenotypic changes along the evolutionary trajectories, we demonstrate how, at the molecular level, the cryptic properties of these evolutionary starting points determine evolvability.

**Figure 1.**
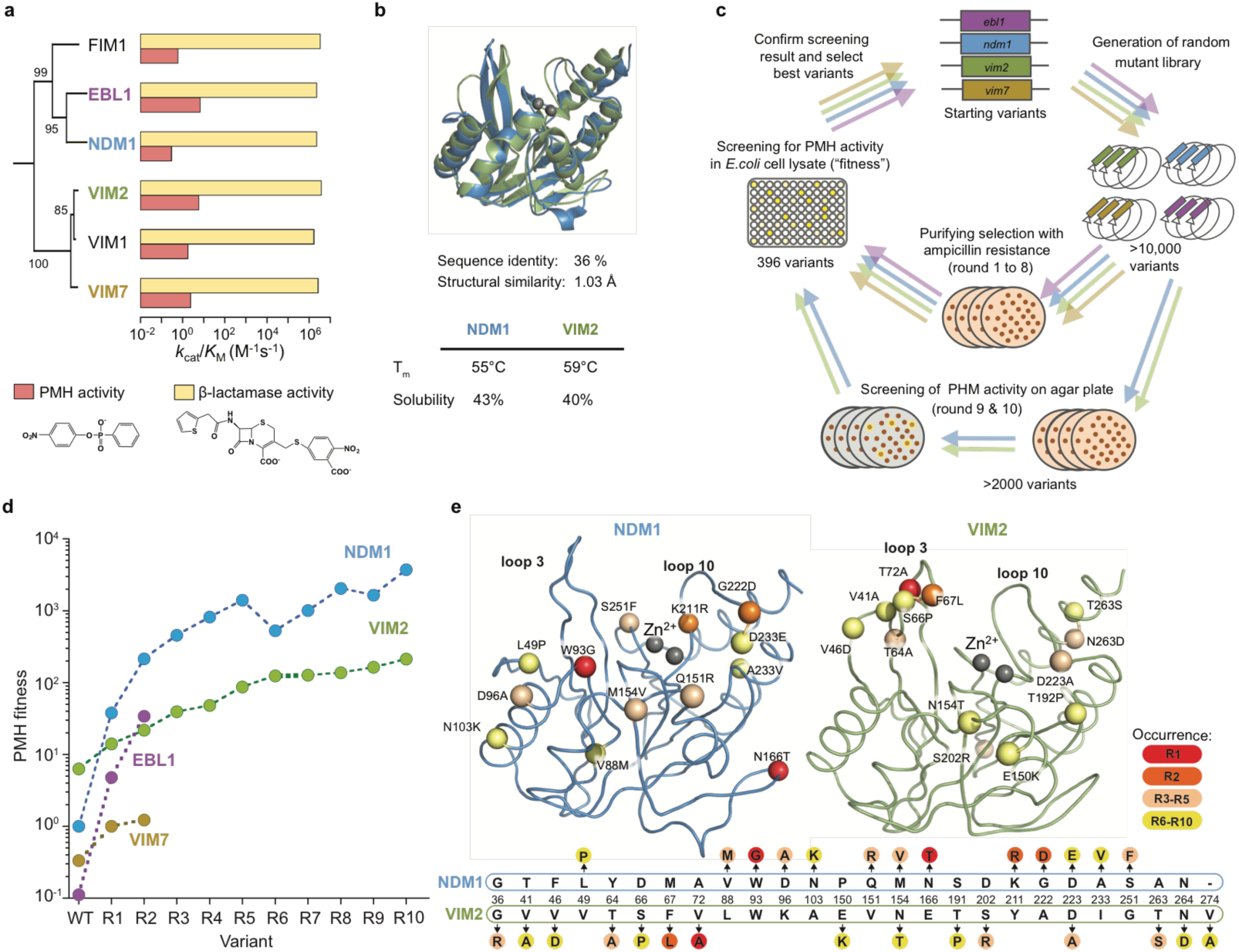
Comparative directed evolution of metallo-β-lactamases (MBLs). **a,** Catalytic efficiencies (*k_cat_*/*K_M_*) of six metallo-β-lactamases (MBLs) for β-lactamase and PMH activities. The phylogenetic relationship is shown with bootstrap values indicated at each node. **b,** Comparison of NDM1 (blue, PDB ID: 3SPU) and VIM2 (green, PDB ID: 1KO3) in overall 3D structure and biophysical properties. **c,** Overview of the comparative directed evolution experiment of MBLs towards PMH activity. **d,** PMH fitness improvements of the directed evolution. PMH fitness is presented as PMH activity in cell lysate relative to NDM1-WT. The activity level of each variant is listed in **Supplementary Table 3**. **e,** The mutations of NDM1 and VIM2 over the ten round of directed evolution. The structural location of the mutations is mapped on the wild-type structures of NDM1 and VIM2 with the C-α atoms of mutated residues shown as spheres (upper panel). Mutations are described on a partial alignment NDM1 and VIM2 sequences (bottom panel). Full alignment of MBLs and mutations of individual variants are presented in **Supplementary Fig. 1** and **Supplementary Table 2**.

## Results

### Comparative directed evolution of four different MBLs

We performed directed evolution using four different MBL orthologs, NDM1, VIM2, VIM7 and EBL1, as starting genotypes (**Fig. 1,** **Supplementary Fig. 1** and **Supplementary Table 1**). The level of the native β-lactamase activity across these enzymes is similar, but their promiscuous PMH activity varies by over 50-fold (**Fig. 1a**). From these starting points, the same directed evolution scheme was applied to improve PMH activity (**Fig. 1c**). Briefly, randomly mutagenized gene pools were transformed into *Escherichia coli*, and purifying selection was used to enrich for functional variants and purge out non-functional variants by plating the library onto agar plates with a low concentration (4 μg/ml) of ampicillin. Colonies were inoculated into 96-well plates (in total 396 variants per round), regrown, lysed and screened for PMH activity. The “enzyme fitness” (or selection criteria) in our directed evolution scheme is defined as the level of PMH activity in *E. coli* cell lysate. The most improved variant(s) were isolated, sequenced and used as templates for the next round of evolution.

We observed significant differences in the degree of fitness improvement among four alternative trajectories over the first two rounds of directed evolution (**Fig. 1d**). Interestingly, improvement in fitness is not correlated to the initial fitness level. For example, the variant with the highest initial activity, VIM2, improved only 4-fold, and was surpassed by the other, initially less fit, variants, NDM1 and EBL1, by 210- and 310-fold improvement, respectively. We continued the directed evolution of VIM2 and NDM1 for an additional eight rounds (**Fig. 1d** and **Supplementary Tables 2**-**3**). Overall, both trajectories demonstrated diminishing returns in their evolution toward new activity, *i.e*. each trajectory eventually reached a plateau^29,30^. Note that the fitness plateaued regardless of purifying selection using β-lactam antibiotic resistance; purifying selection was not employed in the last two rounds (R9 and R10), yet no variant with further improved fitness was isolated (**Fig. 1d**). Besides the similar trend of the trajectory, there were substantial differences in their evolutionary outcomes; NDM1 was initially 4-fold less-fit than VIM2, but its fitness was 28-fold higher by the end of the evolutionary experiment. The NDM1 trajectory improved by 3600-fold, whereas the VIM2 trajectory improved by only 35-fold, resulting in a 100-fold difference in their relative evolvability with respect to PMH fitness. Given that the two WT enzymes were almost identical in terms of their physicochemical properties, protein solubility, stability and structure (**Fig. 1a,b**), the variation in the evolutionary potential that separates these orthologous sequences is substantial.

### Genotypic solutions vary across the evolutionary trajectories

Overall, NDM1 and VIM2 accumulated 13 and 15 mutations respectively. Interestingly, the mutations occurred along each trajectory were entirely distinct (**Fig. 1e**, and **Supplementary Table S2**). Only two mutations occurred at the same position (154 and 223), and these were mutated to different amino acids. For example, the mutations in the early rounds of evolution, which conferred the largest fitness improvements, were W93G, N116T, and K211R in the NDM1 trajectory, and V72A, and F67L for VIM2. In NDM1, most mutations are scattered relatively evenly around the active site; only one mutation (W93G) is located below loop 3, and several are located in and around loop 10 and other parts of the active site. By contrast, in VIM2, six mutations are tightly clustered within or next to loop 3. These results demonstrate that the distinct phenotypic outcomes for the two enzymes result from distinct mutational responses.

### Repeatability and determinism of evolutionary adaptation

An important question that arises from the observation that the two enzymes followed different mutational paths is whether this occurred because of differences in their innate molecular property for evolvability, or if it was due to random chance from experimental variation. To address this, we pursued several lines of evidence that suggest the observed trajectories are largely deterministic (**Fig. 2**). First, we generated and screened two additional libraries from each wild-type enzyme, which repeatedly identified the same mutations that are observed in early rounds of the original directed evolution (W93G in R1 for NDM1 and V72A and F67L in R1 and 2, respectively, for VIM2, **Fig. 2a**). Second, we assessed the epistatic effects of the mutations by introducing the mutations that occurred between R2-R4 into the corresponding wild-type genetic background, and found that the positive effect of later mutations in NDM1 confer a selective advantage only after the fixation of the initial mutations along that trajectory (**Fig. 2b**). Third, we introduced the initial mutation from each trajectory (W93G for NDM1 and V72A for VIM2) into the counterpart enzyme and assayed its effect on fitness, which revealed that each trajectory’s adaptive mutations are incompatible with the other’s. W93G, which increased fitness for NDM1 by 25-fold, caused a 6-fold fitness decrease for VIM2-WT, thus explaining why VIM2 did not acquire this mutation during its evolution (**Fig. 2c**). Introducing other hydrophobic residues (A, V, L and F) at position Trp93 in VIM2 had similar negative effects (**Fig. 2d**). Similarly, V72A, which improved the fitness of VIM2 by around 2-fold, was largely neutral for NDM1 (**Supplementary Fig. 2**, and **Supplementary Table 4**). Taken together, this evidence suggests that intramolecular epistasis results in each genotype only having a limited and unique set of mutations available for adaptation to the PMH activity, which results in mostly repeatable and deterministic evolutionary trajectories.

**Figure 2.**
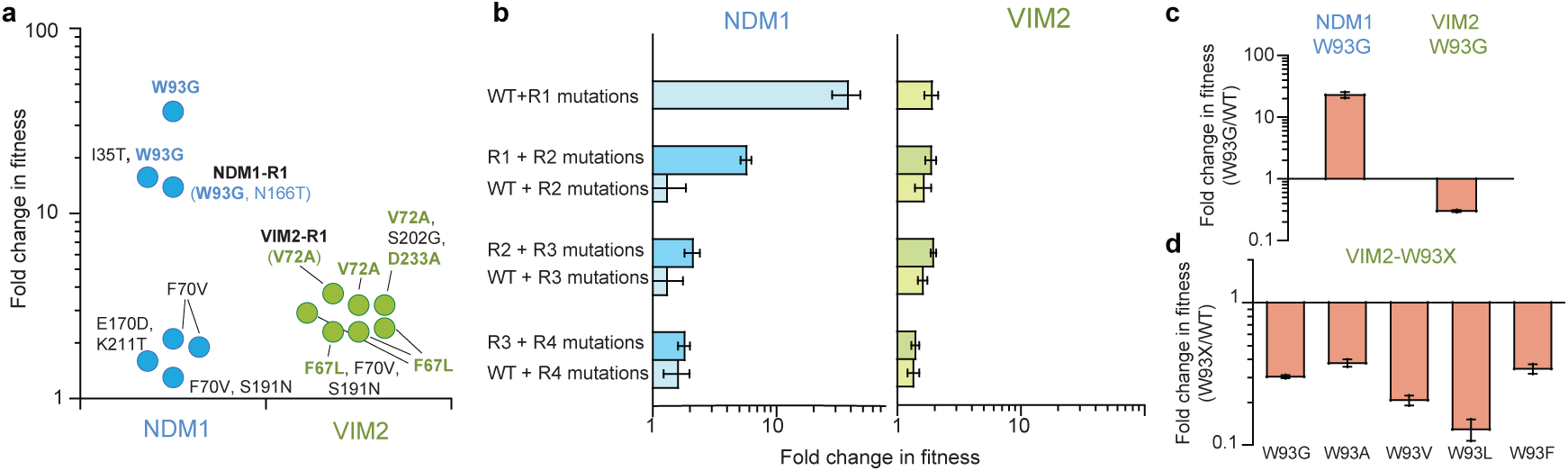
Repeatability of the two evolutionary trajectories. **a,** Additional screening of the wild-type libraries. The improved variants which were isolated from two additional screens of the mutagenized libraries from the wild-type sequences, NDM1-WT and VIM2-WT. Mutations highlighted in colours are the mutations that were isolated in the original directed evolution experiment. **b,** Epistasis analysis of mutations. The mutations occurring in the trajectory of NDM1 and VIM2 were introduced into the respective wild-type sequence and the change in fitness was compared to the ones observed in the trajectory. The errors bars represent the propagated standard deviation from three independent experiments. **c,** Fold change in the PMH fitness of W93G mutants compared to the WT variants. **d,** Fitness effect of introducing various hydrophobic residues at position Trp93 in VIM2.

### Distinct molecular changes underlie the two evolutionary trajectories

The conventional paradigm of protein evolution is dominated by the idea that higher protein stability, or greater soluble expression, promotes protein evolvability, buffering the destabilizing effect of function-altering mutations, allowing a greater number of adaptive mutations to accumulate^23–25^. This model, however, fails to predict the difference in evolvability between NDM1 and VIM2, because their relative stability and solubility are similar (**Fig. 1b**). In order to elucidate the precise molecular changes that enabled each trajectory’s optimization, we measured a range of molecular properties, including catalytic efficiency (*k*_cat_/*K*_M_), solubility, melting temperature (*T*_m_), and oligomeric assembly, over the course of their evolution. The molecular changes that underlie their respective fitness improvements differed substantially (**Figs. 3,** **Supplementary Fig. 3** and **Supplementary Tables 5**-**7**). In NDM1, *k*_cat_/*K*_M_ improved by 20,000-fold from 0.32 to 5,900 M^−1^s^−1^ by R10 (**Figs. 3a-b**). The significant improvement, however, was offset by a loss of protein solubility, which mostly occurred in R1, where *k*_cat_/*K*_M_ increased 300fold, but solubility decreased from 43% to 25% (**Figs. 3c-d**). The level of solubility never recovered, while *k*_cat_/*K*_M_ gradually increased until it reached the plateau observed in R7. By contrast, the *k*_cat_/*K*_M_ of VIM2 stagnated at only a 30-fold increase up to round 6, with the subsequent fitness improvements being due to improvement in solubility from 40% to 70% (**Fig. 3**). Changes in solubility are only weakly correlated to changes in *T*_m_, indicating that other factors such as kinetic stability or protein folding affect the level of soluble protein expression more than thermostability (**Supplementary Fig. 3**). NDM1 retains the same monomeric quaternary structure along its trajectory to NDM1-R10. The monomeric VIM2-WT, however, evolved to exist in an equilibrium between monomer and dimer by VIM2-R10 (**Supplementary Fig. 4a**). We isolated monomeric and dimeric states using size-exclusion chromatography, and determined their respective kinetic parameters. The dimer state is less active than the monomer state for PMH activity, indicating that the dimer formation may be associated with increased in overall solubility along the VIM2 trajectory (**Supplementary Fig. 4b**).

**Figure 3.**
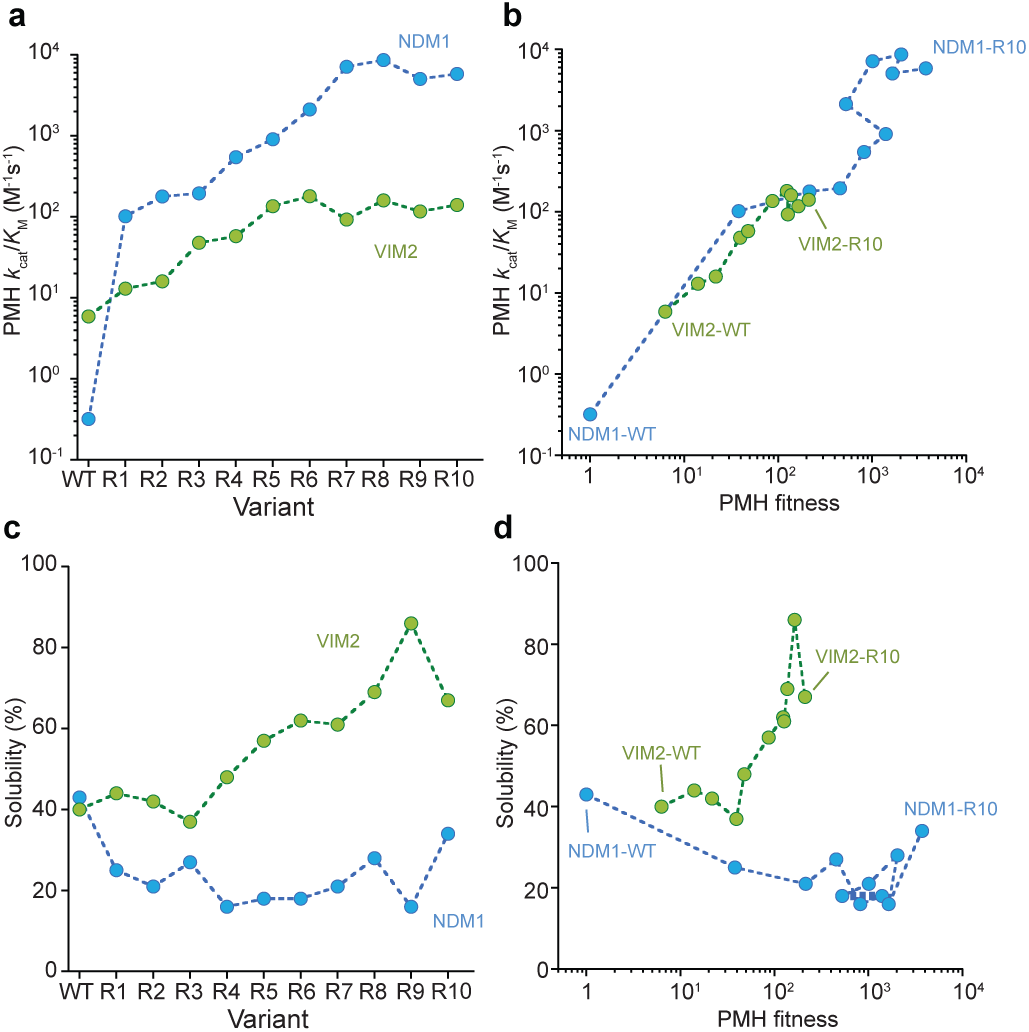
Molecular, phenotypic changes of NDM1 and VIM2 in the evolutionary trajectory. **a,** Catalytic efficiencies (*k_cat_*/*K_M_*) of purified variants for PMH activity. Individual catalytic parameters are listed in **Supplementary Tables 5**-**6**. **b,** Correlation between fitness and catalytic efficiency (k_cat_/K_M_) along the evolutionary trajectories. **c,** Changes in solubility in the evolutionary trajectory. SDS-PAGE of each variant was presented in **Supplementary Fig. 3.d**, Correlation between fitness and solubility along the evolutionary trajectories.

Altogether, our results confirm that differences in protein stability cannot explain the differences in evolvability between the two enzymes. VIM2 variants consistently exhibit higher stability and solubility throughout the trajectory, yet the improvements in *k*_cat_/*K*_M_ and fitness are far lower than that of NDM1. Moreover, the distinct phenotypic solutions further highlight the qualitatively different evolutionary processes that led to each trajectory’s fitness plateau. Interestingly, the genetic differences in the evolutionary end points can create additional cryptic variation that becomes apparent when the environment is changed: when the enzymes are expressed at 37°C (rather that the 30°C at which they evolved PMH activity) NDM1 variants are significantly less fit because of their lower soluble expression, whereas VIM2 variants maintain similar fitness levels even at the higher temperature (**Supplementary Fig. 5**).

### Structural adaptation between the evolutionary trajectories

Having established that protein stability does not constrain the evolvability of the enzymes, we sought a molecular explanation by solving the crystal structures of the R10 variants for both trajectories, allowing to us compare them with the previously published wild-type structures (**Fig. 4** and **Supplementary Table 8**)^31,32^. For NDM1-R10 we obtained crystal structures of the apo-enzyme and a complex with the phenylphosphonate product bound in the active site after *in crystallo* substrate turnover. Additionally, we conducted molecular dynamics (MD) simulations in complex with the PMH substrate (*p*-nitrophenyl phenylphosphonate, PPP). We identified three main structural adaptations from NDM1-WT that underlies the 20,000-fold improvement in PMH activity. First, W93G removes the steric hindrance between the side chain of Trp93 and the substrate, and generates a complementary pocket for the phenyl group below loop 3 (**Figs. 4b** and **S6**). Second, there is a displacement of loop 3 (Trp93 is located near the base of this loop) inward by ~6 Å, which allows for improved π-π stacking interactions between Phe70 and the *p*-nitrophenol-leaving group (**Figs. 4c** and **Supplementary Fig. 6**). Third, loop 10 is repositioned *via* reorganization of the local hydrogen bond network (largely by K211R and G222D), allowing it to interact with the leaving group of the substrate (**Fig. 4c**). For VIM2, we were only able to crystallize the dimer fraction of VIM2-R10. This dimer reveals an unprecedented structural rearrangement: a full half of the structure is symmetrically domain-swapped between two subunits (**Figs. 4d** and **Supplementary Fig. 6**). Besides the domain swapping, the major structural arrangement between VIM2-WT and VIM2-R10 involves the reorganization loop 3, as loop 3 is disordered in VIM2-R10, which is caused by six mutations that occurred within and next to loop3 (**Figs. 1e** and **4d**). Four of these six these loop 3 mutations were accumulated by R5, and thus we speculate that these mutations and the loop rearrangement was the major cause of 30-fold increase in PMH activity in VIM2. Taken together, the results further emphasize that the two starting enzymes responded to the same selection pressure by substantially different molecular changes.

**Figure 4.**
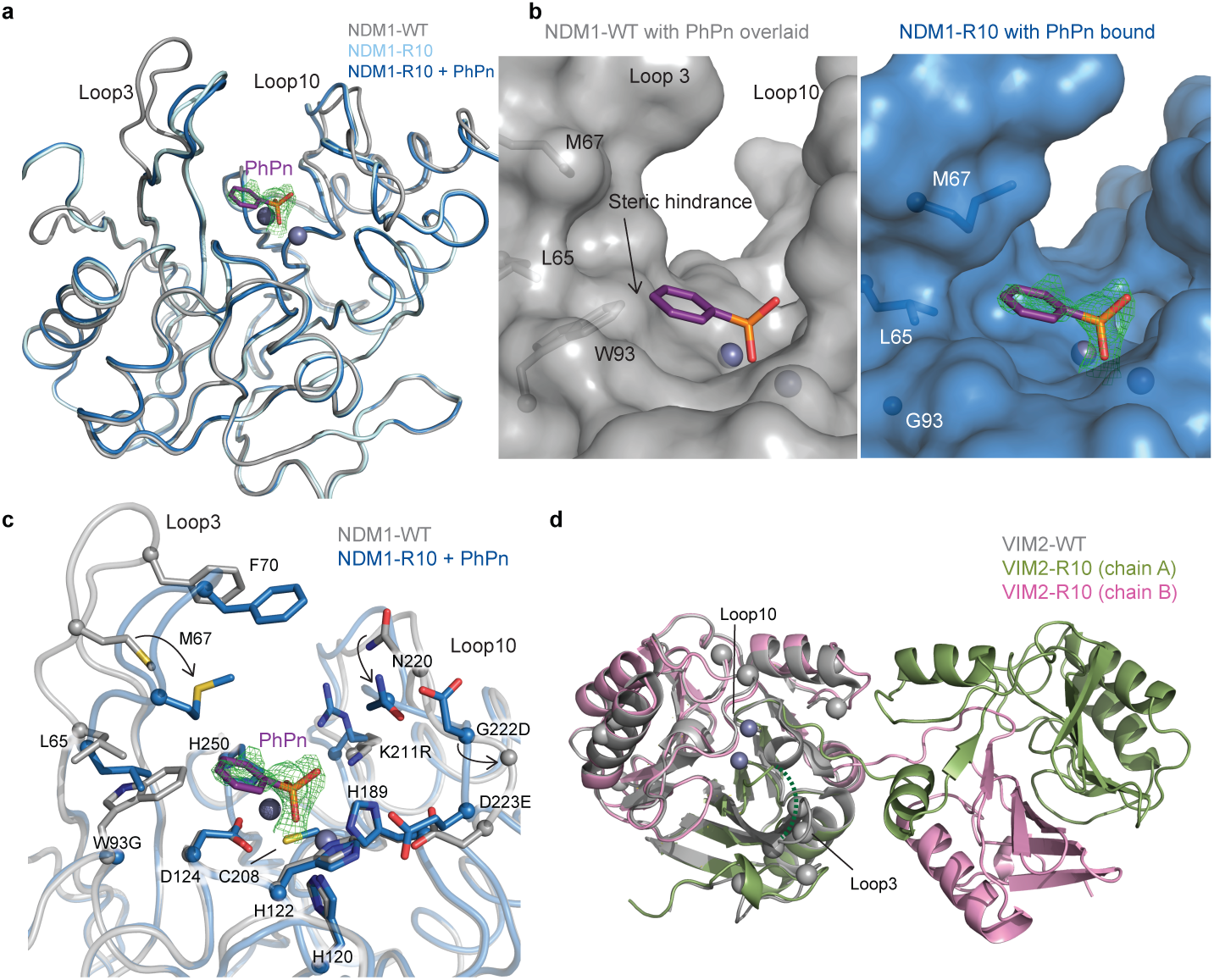
The structural basis for improved PMH activity. **a,** Structural overlay of NDM1-WT (grey, PDB ID: 3SPU) and NDM1-R10 in the apo (cyan, PDB ID: 5JQJ) and the PMH product, phenyl-phosphate (magenta sticks) complexed form (blue, PDB ID: 5K4M). The active site metal ions are shown as spheres. **b,** Surface views of the active site of NDM1-WT and NDM1-R10 with the PMH product. The product in the NDM1-WT structure is replaced based on the NDM1-R10 structure. The mFo-dFc omit electron density of the pheyl-phosphate is shown (green mesh), contoured at 2.5σ. **c,** Comparison of active site residues between NDM1-WT and NDM1-R10. **d,** Structural overlay of monomeric VIM2-WT (grey, PDB ID: 1KO3) and domain swapped-dimeric VIM2-R10 (green and pink, PDB-ID: 6BM9). The disordered loop 3 in VIM-R10 is indicated as a dashed line. The mutations accumulated during the VIM2 trajectory were depicted as light grey spheres. The mFo-Fc omit electron density of the structure is shown in **Supplementary Fig. 7**.

### Molecular basis for the mutational incompatibility of the key mutation W93G

Finally, we investigated the molecular basis for a key mutation that differentiates the two trajectories: W93G. This mutation caused a 100-fold increase in the catalytic activity of NDM1, but when introduced into VIM2, it reduces PMH activity by 10-fold (**Supplementary Fig. 2**). We performed and compared MD simulations of NDM1-WT, VIM2-WT and models of NDM1-W93G, and VIM2-W93G in the presence of the PMH substrate (**Figs. 5,** **Supplementary Fig. 8** and **Supplementary Tables 9****-10**). NDM1-W93G showed similar structural adaptations as observed in NDM1-R10 (described above); W93G eliminates the steric hindrance with the substrate, and allows loop 3 to shift inward to promote complementary interactions with the substrate, but without the rearrangement of loop 10, which presumably occurs later in that evolutionary trajectory. By contrast, in VIM2-WT, Trp93 adopts a different orientation, avoiding steric hindrance and instead promotes complementary interactions with the phenyl ring of the substrate. Consequently, W93G in VIM2 removes beneficial substrate-enzyme interactions, thus causing a deleterious effect on PMH activity. Similarly, unlike in NDM1, in VIM2-W93G, loop 3 does not form complementary interactions with the substrate, which is consistent with the observation that loop 3 is extensively mutated and reorganized by other mutations later in the VIM2 trajectory. By examining these crystal structures, we found that the “second-shell” residues of Trp93 cause different orientations of the indole sidechain. In NDM1, Leu65, Gln123 and Asp124 constrain Trp93 to point into the active site (**Fig 5d**). In VIM2, however, the second shell residues (Gln65, Asp123, and Asp124) differ, resulting in Trp93 being stabilized in an alternative conformation (**Fig. 5e**). Thus, remote and seemingly neutral sequence variation between the enzymes has a substantial impact on the mutagenesis of a key active site residue.

**Figure 5.**
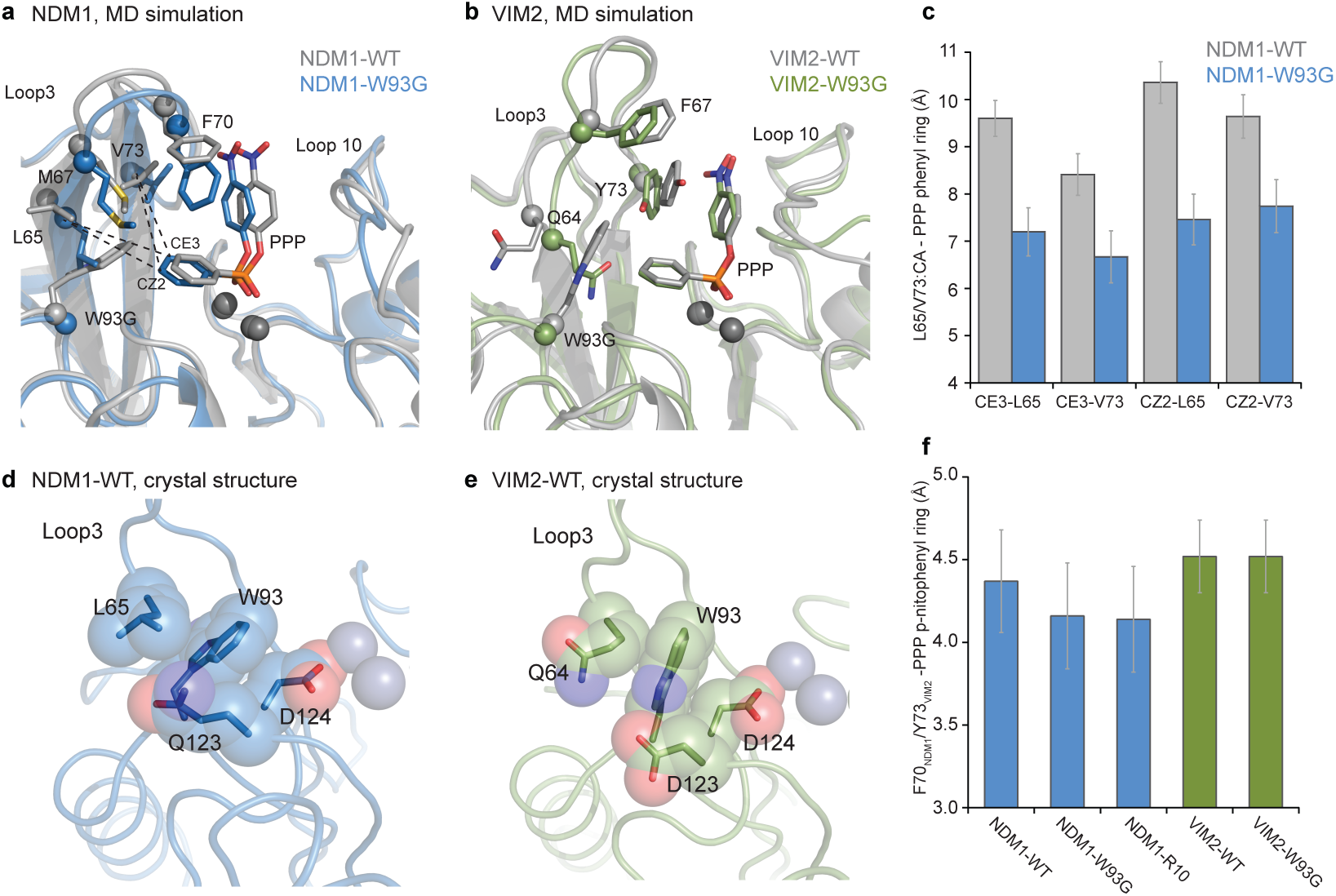
Comparisons of structural changes by the mutation W93G. Representative structures from MD simulations of **(a)** NDM1-WT (grey) and NDM1-W93G (blue), and **(b)** VIM2-WT (grey) and VIM2-W93G (green). **c,** Changes in the hydrophobic interactions of the phenyl ring of the PMH substrate upon the W93G mutation observed in the MD simulations of NDM1, quantified through average distances between CE3 and CZ2 atoms of the phenyl ring of PMH and the alpha carbon atoms of residues Leu65 and Val73. **d,** The position of Trp93 and second shell residues in the NDM1-WT crystal structure (PDB-ID: 3PSU). **e,** The position of Trp93 and second shell residues in VIM2-WT crystal structure (PDB-ID: 1KO3). **f,** The average distance between the *p*-nitrophenyl ring of the substrate and side chain rings of Phe70 in NDM1 and Tyr73 in VIM2, when the two rings form a *π*-*π* interaction during the MD simulations.

We expanded our mutational analysis of W93G and V72A (the first mutation in the VIM2 trajectory) to four other orthologous enzymes (EBL1, FIM1, VIM1, VIM7) and found that their effects for both PMH and β-lactamase activities consistently vary significantly, even between orthologs with high sequence identity (**Supplementary Fig. 2**). For example, in VIM7, which is 80% identical to VIM2, W93G caused a 3-fold increase in activity, despite the same mutation causing a 10-fold decrease in VIM2. Taken together, cryptic and subtle differences in sequence and structure, can influence the conformation of a key active site residue, causing an approximately 3000-fold difference in the phenotypic effect of a mutation, and thereby leading to distinct evolutionary outcomes among orthologous enzymes.

## Discussion

The great diversity of protein functions, and many contemporary examples of proteins that have promptly adapted to changing environments suggest a remarkable degree of evolvability for biological molecules^33,34^.These successful cases of adaptation, however, may also obscure a wealth of cases where proteins failed, or were limited, in their evolution of a new function. Our observations highlight that not all enzymes are equally evolvable, and that seemingly innocuous genetic variation can result in significant consequences for a protein’s ability to evolve a new function. Thus, this work elaborates on the known role of cryptic genetic variation in generating diverse, hidden “pre-adapted” properties^5–7^, extending this view to encompass the adaptive evolutionary potential of individual genes, with meaningful implications for the evolvability of organisms and populations as well.

Our observations, when combined with those of others, indicate that the evolvability of proteins is generally and profoundly rooted in their genetic variation. Examples from nature and the laboratory have shown that independent evolution trials from a single genotype often follow very similar genetic and evolutionary trajectories, suggesting that evolution is largely deterministic from a particular genetic starting point^35–40^. On the other hand, the prevalence of epistasis causes distinct mutational effects amongst potential adaptive mutation(s) for otherwise phenotypically similar orthologs^14–17^, suggesting that genetic variation epistatically “restricts” and/or “permits” the accessibility of certain adaptive mutations ^12,41,42^. The successful evolution of new protein functions may therefore rely on genetic drift to explore the sequence space, and generate diverse genotypes with differently evolvable genotypes^16^. The genetic diversity that these neutral drifts generate thereby provide a foundation for the response of a population to new selection pressures^9,41^. One complicating aspect of this model is the impact of temporal and spatial occurrences of selection pressure on the emergence of diverse genotypes. It is possible that this model explains a commonly observed phenomenon in bacterial adaptation, *e.g*., drug-resistance and xenobiotic degradation^43,44^, in which typically there are only a small number of genotypes in a larger population that emerge to confer new functions, after which point these successfully adapted genes quickly disseminate to other bacteria *via* horizontal gene transfer.

Our observations suggest that the conventional paradigm, where protein stability is the dominant factor in determining evolvability^23–25^, cannot account for variation in evolutionary potential between the MBL enzymes toward PMH activity. Instead, we found that cryptic and subtle molecular differences have a far greater impact on enzyme evolvability. Moreover, the initial fitness and phenotypes of the enzymes does not necessarily provide a good indicator for their eventual evolutionary outcomes. Thus, adaptive evolutionary potential can be truly “cryptic” and only apparent after evolution happens, and further it is extremely difficult to predict.

These results also have profound implications for protein design, engineering and laboratory evolution: protein engineers overwhelmingly choose a single starting genotype based on the availability of biochemical and structural information (often the highest initial activity), and much effort has been devoted to develop technologies to overcome evolutionary dead-ends^45–47^. Our observations suggest that it would be more effective to explore diverse genotypes and identify the most evolvable starting sequences to successfully obtain an optimized functional protein. The biophysical rationalization of the cryptic properties of MBL enzymes described here contribute to the ambitious goal of understanding the molecular mechanisms behind neutral genetic variation and evolvability that we observe in nature, in such a way as to allow us to predict evolutionary pathways and understand how to acquire better biological molecules.

## Acknowledgements

We thank Dan S. Tawfik, Amir Aharoni, Joelle Pelletier and the members of the Tokuriki lab for comments on the manuscript. Natural Sciences and Engineering Research Council of Canada (NSERC) Discovery Grant (RGPIN 418262-12 and RGPIN 2017-04909), and Canadian Institute of Health Research (CIHR) Foundation Grant to N.T.. N.T. is a CIHR new investigator and a Michael Smith Foundation of Health Research (MSFHR) career investigator. We also thank the Knut and Alice Wallenberg and Wenner-Gren Foundations for fellowships to S.C.L.K. and A.P. respectively, as well as the Swedish National Infrastructure for Computing (SNAC) for supercomputing resources.

## Author contributions

F.B. and N.T. conceived and designed this study. F.B. and G.Y. performed experimental evolution, enzymatic assay and mutational analysis. N.H., P.D.C., C.J.J. collected structural data. A.P., A.B. and S.C.L.K. designed and conducted MD simulations. F.B. and N.T. wrote the paper with input from all authors.

---

**Supplementary Figure 1.**
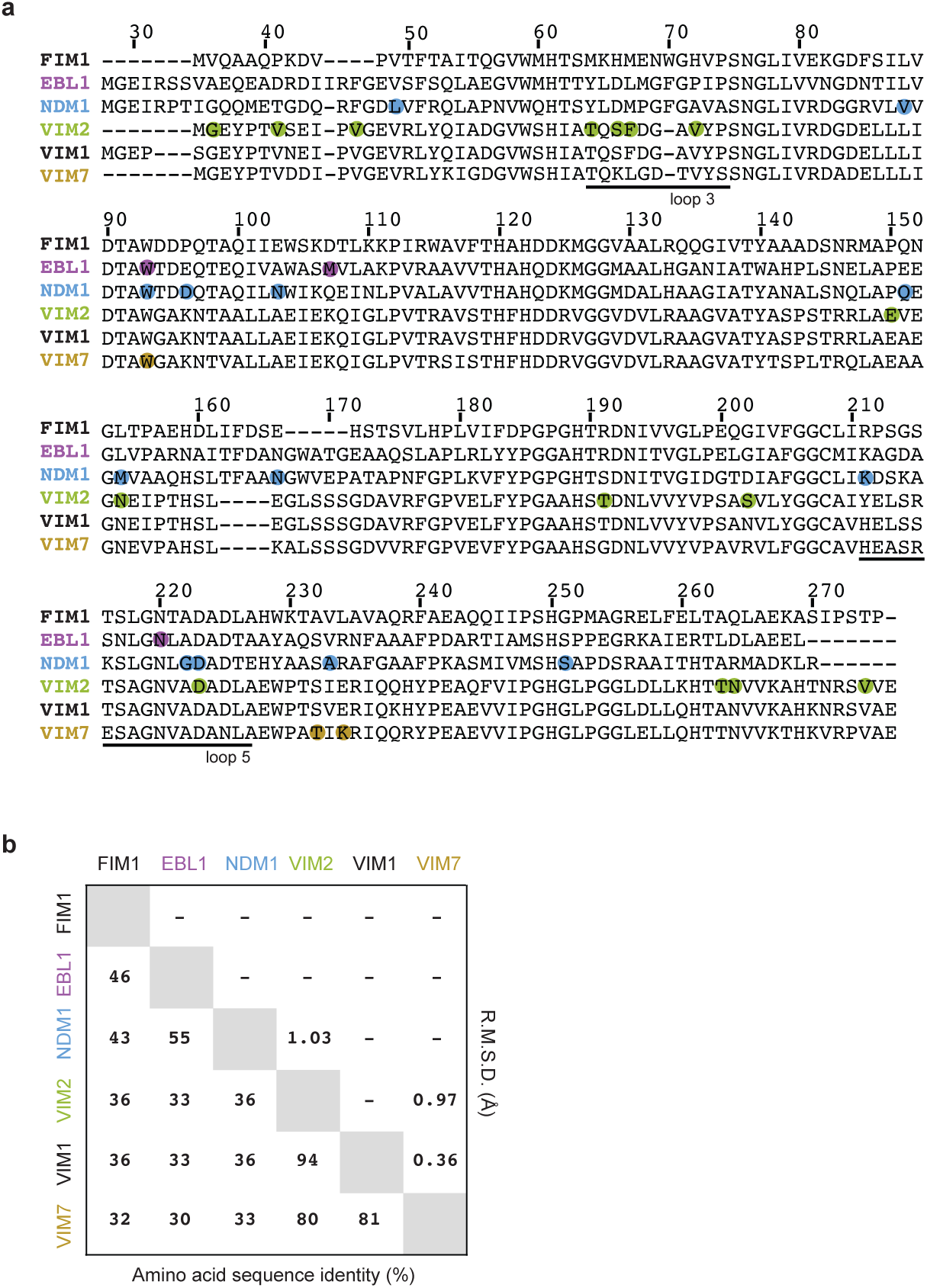
Sequence comparison between metallo-β-lactamases. **a**, Sequence alignment of selected metallo-β-lactamases. Residue numbering is based on the NDM1 sequence (PDB-ID 3spu). Positions that were mutated and selected during the directed evolution for PMH activities are highlighted in color. Actual mutations are listed in **Supplementary Table 2.b**, Sequence identity and structure similarity among selected B1 β-lactamases. A multiple sequence alignment of the nine B1 β-lactamases using ClustalW2 (standard parameters), which was then used to calculate the pairwise sequence identities using the web based program SIAS (hcp://imed.med.ucm.es/Tools/sias.html) with gaps taken into account. To determine pairwise structural similarity, we computed the root mean standard deviation (RMSD) between all structure pairs using the align command in PyMOL.

**Supplementary Figure 2.**
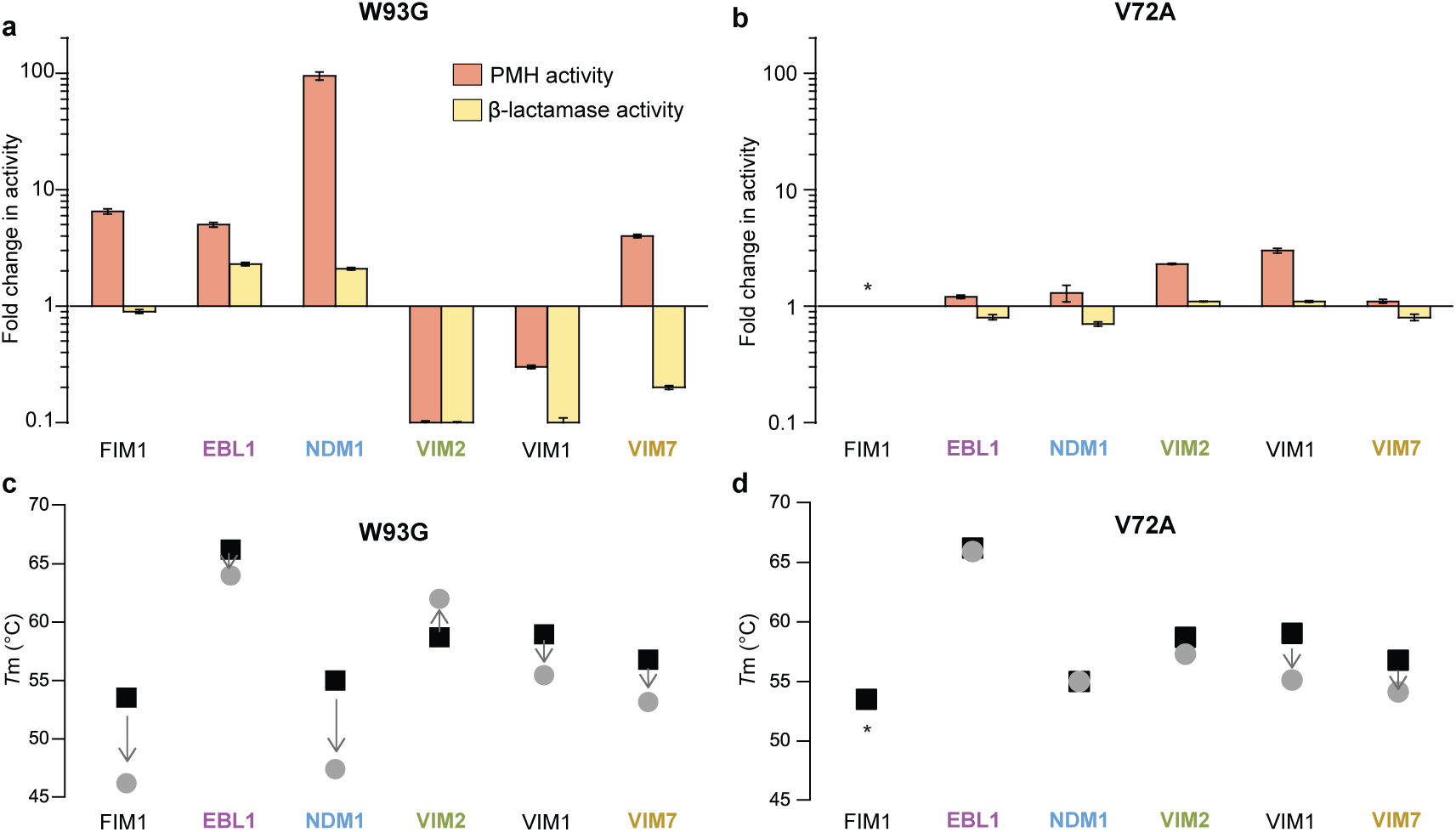
Differential effects of mutations (W93G and V72A) on six metallo-β-lactamases. Fold change in PMH and β-lactamase activity of variants. (**a**) W93G and (**b**) V72A variants compared to the wild type enzyme. Activity levels of purified enzymes were measured at single enzyme (1 μM for PMH and 1 nM for β-lactamase activity) and substrate (500 μM for PMH and 100 μM for β-lactamase activity) concentrations. Errors bars represent the standard deviation calculated from three measurements. Changes in melting temperature (*T*_m_) of (**c**) W93G and (**d**) V72A. Black squares denote the wild-type enzyme, and grey circles denote the mutant. Arrows indicate the change of melting temperature by the mutation. The melting temperature was calculated from the midpoint of the thermal denaturation curve of purified proteins. Asterisks indicate that the fold change in activity could not be determined, because one of the variants was not soluble. The catalytic activities and *T*_m_ of each mutant are listed in **Supplementary Table 4**.

**Supplementary Figure 3.**
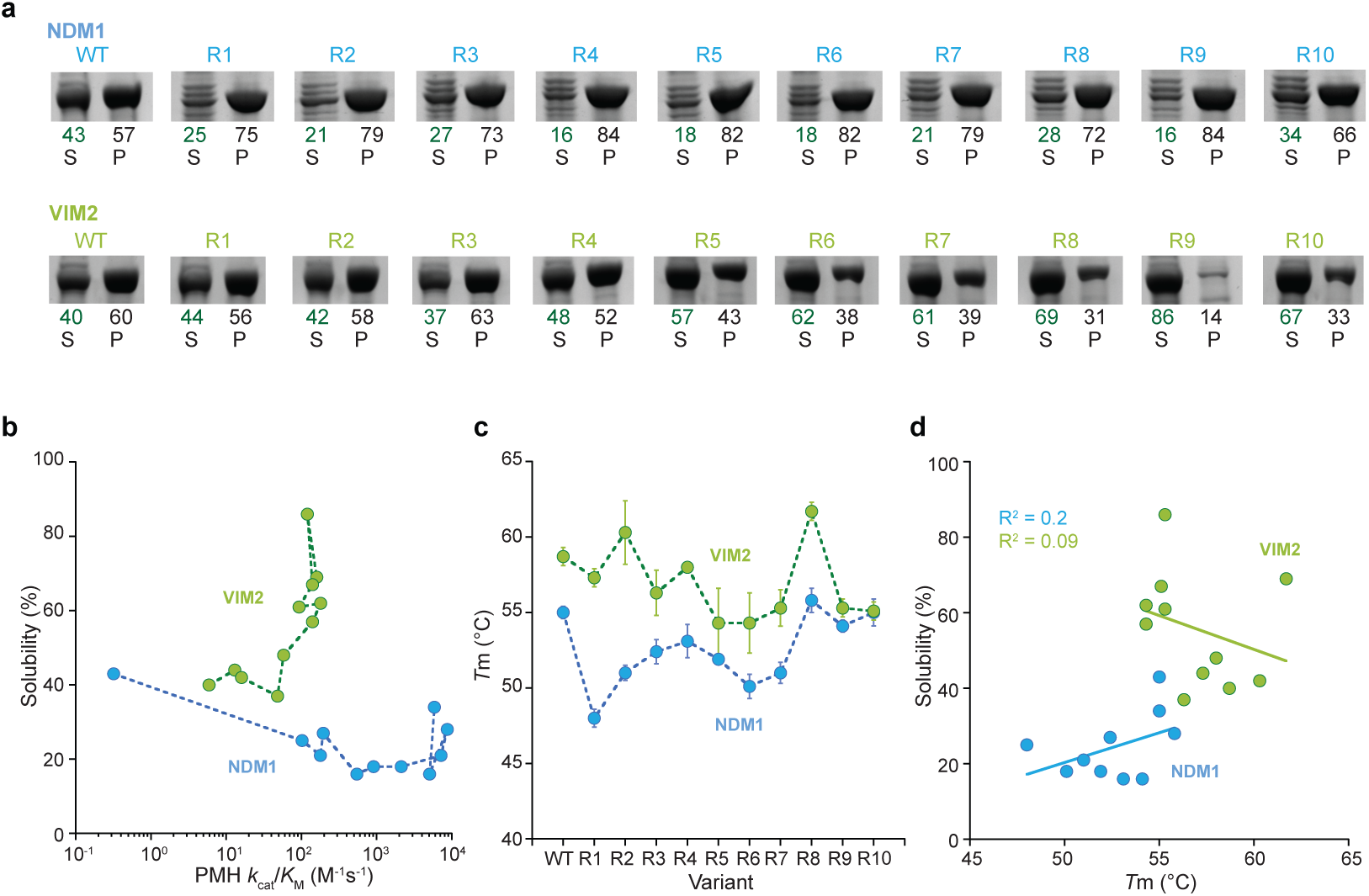
Solubility and melting temeparature of variants. **a**, SDS-PAGE images of protein expression in the supernatant (S) and pellet (P) fractions of the cell lysate. The numbers below indicate the percentage of protein in the soluble and insoluble fraction, which is calculated from the relative intensities of the supernatant and pellet bands. **b**, Correlation between solubility and catalytic efficiency along the evolutionary trajectories. **c**, Melting temperature (*T*_m_) of all variants along the evolutionary trajectory as calculated from the midpoint of the denaturation curve in a thermal shift assay. Error bars represent standard deviation from three independent assays. **d**, Correlation between solubility and melting temperature (*T*_m_) of NDM1 and VIM2 variants.

**Supplementary Figure 4.**
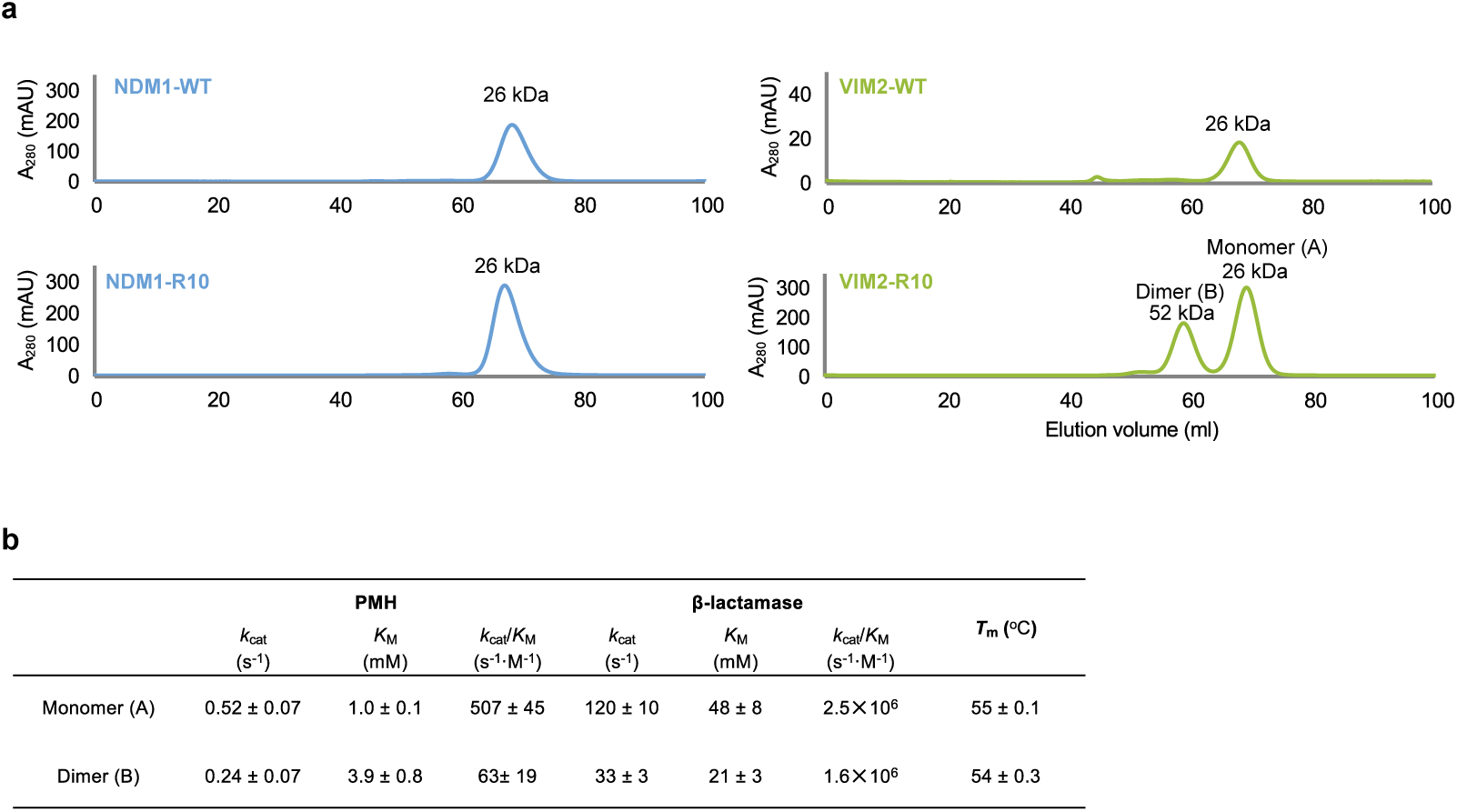
Oligomerization states of NDM1 and VIM2.**a**, Size-exclusion chromatography (SEC) elution profile of NDM1-WT, NDM1-R10, VIM2-WT and VIM2-R10 (Superdex 75, GE Healthcare). Protein sizes were identified based on the calibration curve on the manufacturer’s instruction (GE Healthcare) for gel filtration calibration kits LMW (low molecular weight). VIM-R10 showed two peaks that correnpond to monomeric and dimeric states. **b**, kinetic parameters and melting temperature (*T*_m_) of monomer and dimer states of VIM2-R10.

**Supplementary Figure 5.**
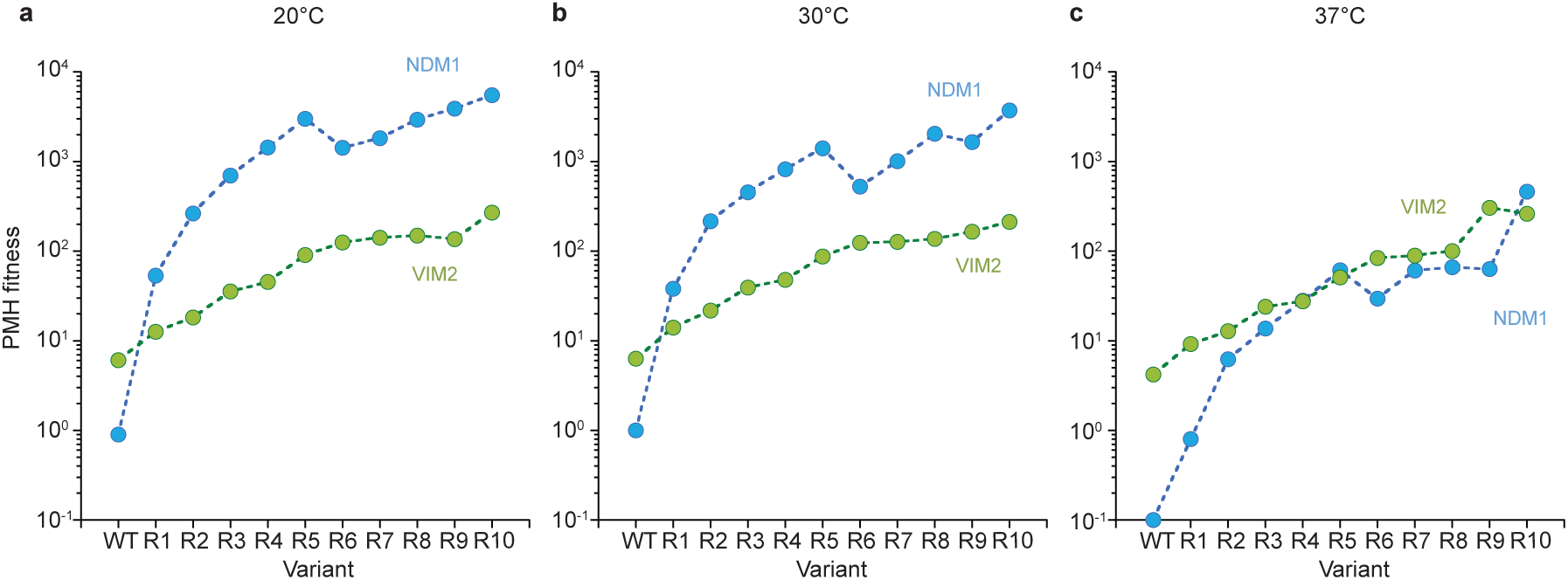
Effect of expression temperature on cell lysate PMH activities. Variants of NDM1 (blue) and VIM2 (green) were expressed at (**a**) 20°C, ( **b**) 30°C and ( **c**) 37°C. Cell lysate preparation and PMH fitness (cell lysate activity) measurement were performed identically for all expression temperatures.

**Supplementary Figure 6.**
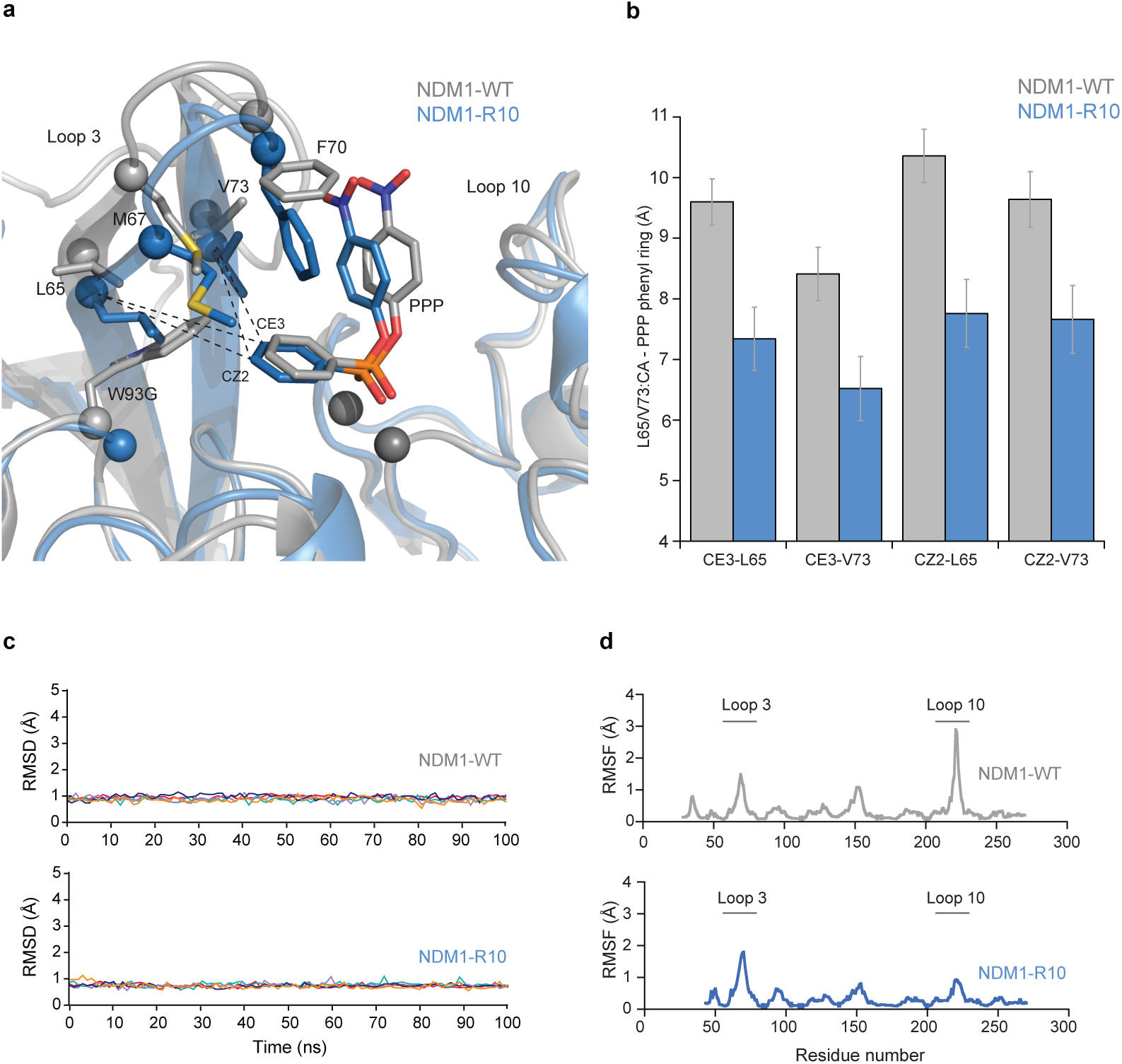
The molecular dynamics (MD) simulations of NDM1-WT and NDM1-R10. **a**, Overlay of representative structures from the MD simulations of NDM1-WT (grey), NDM1-W93G (blue) in complex with the PPP substrate. **b**, The average distance between the substrate and active site residues Leu65 and Val73 during the MD simulation of NDM1-WT and R10. **c**, Root-mean square deviations (RMSD, Å) of all C-α atoms from the MD simulations of the NDM1-WT and NDM1-R10 in complex with PPP. The relevant enzyme variants are indicated on each panel. The data is presented individually for five independent 100 ns trajectories in each system, and for clarity the data was plotted using cspline smoothing function implemented in Gnuplot. **d**, Root-mean square fluctuations (RMSF, Å) of C-α atoms calculated from the last 50 ns of all five 100 ns simulations of NDM1-WT and NDM1-R10 in complex with the PPP substrate.

**Supplementary Figure 7.**
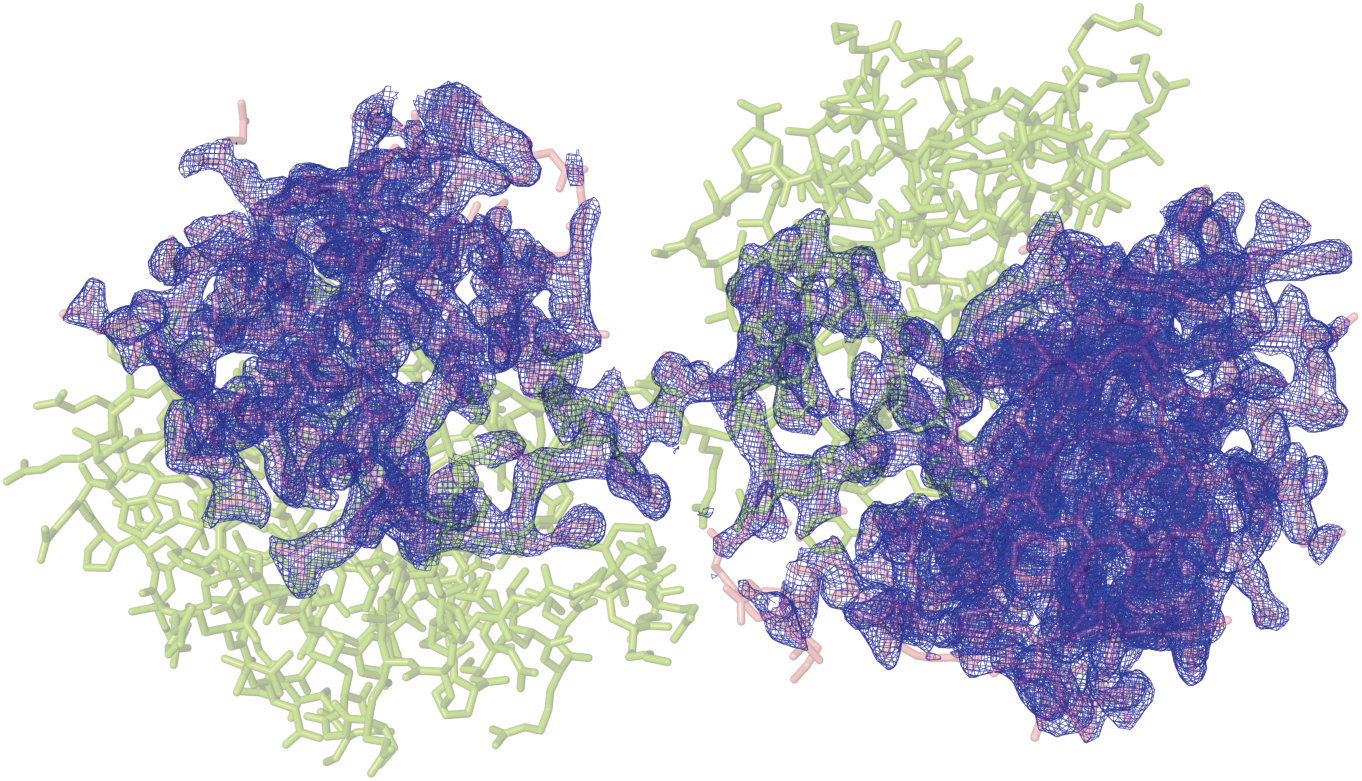
Structure of the domain swapped-dimeric VIM2-R10. A stick representation of the structure of the domain swapped-dimeric VIM2-R10 (PDB-ID: 6BM9). Chain A is highlighted as green, and chain B as pink. The mFo-dFc omit electron density of chain B is shown (blue mesh), contoured at 2.5σ.

**Supplementary Figure 8.**
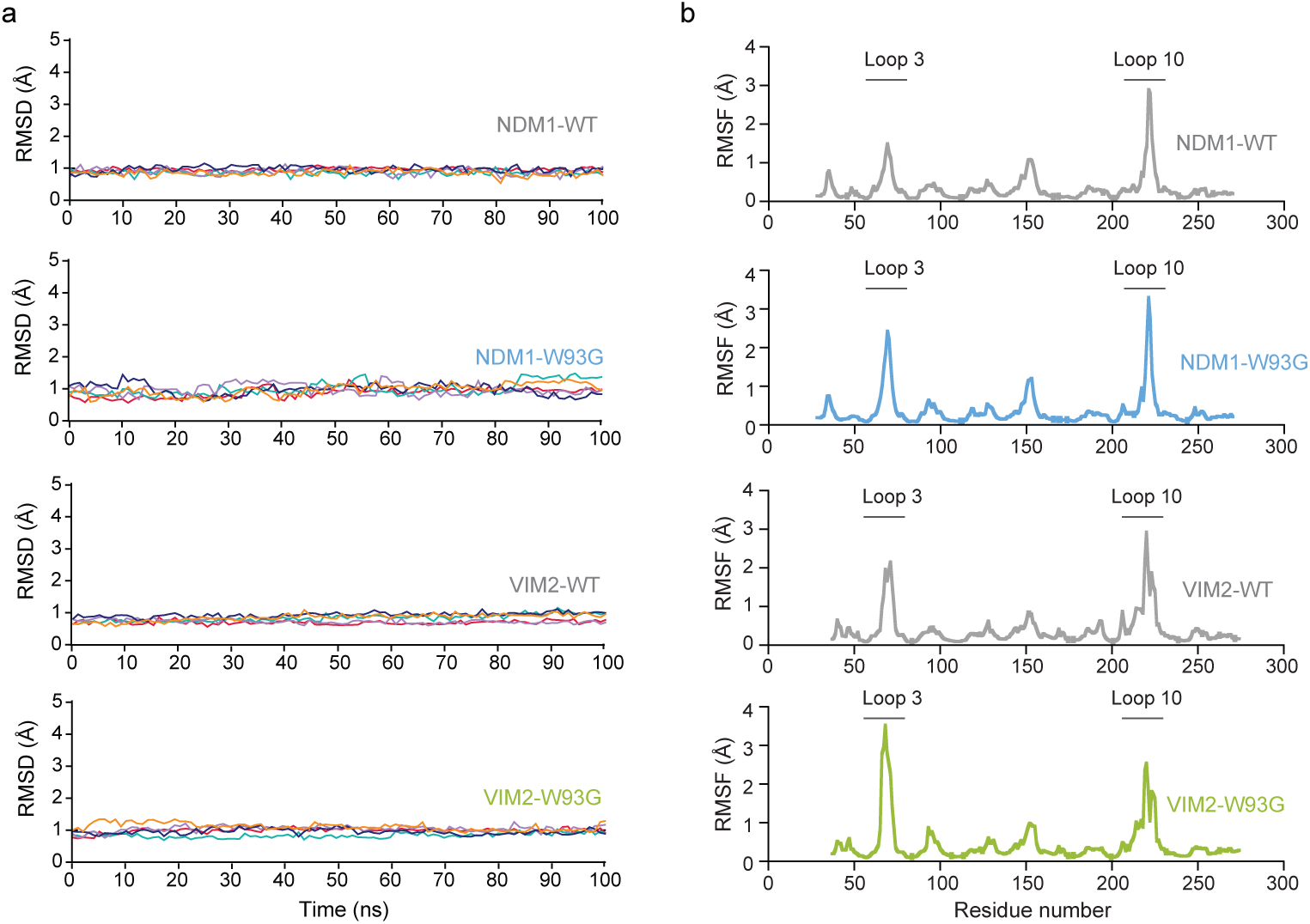
The molecular dynamics (MD) simulations of NDM1-WT, NDM1-W93G, VIM2-WT, and VIM2-W93G. **a**, Root-mean square deviations (RMSD, Å) of all C-α atoms from the MD simulations of NDM1-WT NDM1-W93, VIM2-WT and VIM2-W93G in complex with PPP. The relevant enzyme variants are indicated on each panel. The data is presented individually for five independent 100 ns trajectories in each system, and for clarity the data was plotted using cspline smoothing function implemented in Gnuplot. **b**, Root-mean square fluctuations (RMSF, Å) of C-α atoms during the MD simulations. The data was calculated from the last 50 ns of all five 100 ns simulations of each of the four enzyme-PPP complexes (specific variant indicated in each panel).

## Material and Methods

### Generation of mutagenized library

Random mutant libraries were generated with error-prone PCR using nucleotide analogues (8-Oxo-2′-deoxyguanosine-52′-Triphosphate (8-oxo-dGTP) and 22′-Deoxy-P-nucleoside-52′-Triphosphate (dPTP); TriLink). Two independent PCR reactions were prepared, one with 8-oxo-dGTP and one with dPTP. Each 50 μl reaction contained 1 × GoTaq Buffer (Promega), 3 μM MgCl_2_, 1 ng template DNA, 1 μM of primers (forward (T7 promoter): taatacgactcactataggg; reverse (T7 terminator): gctagttattgctcagcgg), 0.25 mM dNTPs, 1.25 U GoTaq DNA polymerase (Promega) and either 100 μM 8-oxo-dGTP or 1 μM dPTP. Cycling conditions: Initial denaturation at 95°C for 2 minutes followed by 20 cycles of denaturation (30 seconds, 95°C), annealing (60 seconds, 58°C) and extension (70 seconds, 72°C) and a final extension step at 72°C for 5 minutes. Subsequently, each PCR was treated with *Dpn* I (Fermentas) for 1 h at 37°C to digest the template DNA. PCR products were purified using the Cycle Pure PCR purification kit (E.N.Z.A) and further amplified with a 2 × Master mix of Econo TAQ DNA polymerase (Lucigen) using 10 ng of the template from each initial PCR and the same primers at 1 μM in a 50 μl reaction volume. Cycling conditions: Initial denaturation at 95°C for 2 minutes followed by 30 cycles of denaturation (30 seconds, 95°C), annealing (20 seconds, 58°C) and extension (70 seconds, 72°C) and a final extension step at 72°C for 2 minutes. The PCR products were purified and cloned as described above. The protocol yielded 1-2 amino acid substitutions per gene.

### Generation of DNA shuffling libraries

The staggered extension process (StEP) protocol was used to recombine equally improved mutants^48^. Plasmids of variants were mixed in equimolar amounts to 500 ng of total DNA and used as a template for the StEP reaction. Cycling conditions: Initial denaturation at 95°C for 5 minutes followed by 100 cycles 95°C for 30 s followed by 58°C for 5 s. PCR products were purified using the Cycle Pure PCR purification kit and further amplified with a 2 × Master mix of Econo TAQ DNA polymerase. Libraries were cloned into pET29(b) as described above.

### Site-directed mutagenesis

Single-point mutant variants were constructed by site-directed mutagenesis as described in the QuikChange Site-Directed Mutagenesis manual (Agilent) using specific primers. All variants contained only the desired mutation, which was confirmed by DNA sequencing.

### Pre-screen on agar plates

Libraries in pET29-pMBP were electroporated into *E. coli* BL21 (DE3) cells and incubated for 1 h at 37°C prior to plating. For the low antibiotic prescreen, the transformants were plated on agar plates (150 mm diameter) containing 4 μg/ml ampicillin, 0.1 mM IPTG, 200 μM ZnCl_2_ and 40 μg/ml kanamycin, yielding >500 colonies. The minimum inhibitory concentration (MIC) of ampicillin for *E.coli* cells expressing NDM1 and VIM2 is 256 μg/ml, whereas for the *E.coli* cells alone it is <2 μg/ml. Subsequently, surviving colonies were directly picked from plates for rescreening in 96-well plates. For the direct PMH prescreen, transformation reactions were plated on six agar plates (150 mm diameter) containing 40 μg/ml kanamycin, such that each plate contained between 400-2000 colonies. Colonies were replicated onto nitrocellulose membrane (BioTrace NT Pure Nitrocellulose Transfer Membrane 0.2 μm, PALL Life Sciences), which was then placed onto LB agar plates containing 1 mM IPTG, 200 μM ZnCl_2_ and 40 μg/ml kanamycin for overnight protein expression at room temperature. After expression, the membrane was placed into an empty petri dish and the cells were lysed by alternating incubations at −20°C and 37°C three times for 10 min each. To assay activity, 25 ml of 0.5% Agarose in 50 mM Tris-HCl buffer pH 7.5 containing 200 μM ZnCl_2_ and 250 μM *p*-nitrophenyl phenylphosphonate (Sigma) was poured onto the membrane. Colonies with active enzymes developed a yellow colour due to the hydrolyzed substrate. The most active colonies (~200 variants) were directly picked from plates for screening in 96-well plates.

### Cell lysate activity screen in 96-well plates

To test the fitness and solubility of the variants, individual wells of a 96-well plate containing 400 μl of LB media supplemented with 40 μg/ml kanamycin were inoculated with 20 μl of overnight culture and incubated at 30°C for 3 hours. Protein expression was induced by adding IPTG to a final concentration of 1 mM and further incubation at 30°C (20°C and 37°C for testing temperature effect on expression) for 3 hours. Cells were harvested by centrifugation at 4,000 × g for 10 min and pellets were frozen −80°C for at least 30 min. For lysis, cell pellets were resuspended in lysis buffer (50 mM Tris-HCl pH 7.5, 100 mM NaCl, 200 μM ZnCl_2_, containing 0. 1 % Triton X-100, 100 μg/ml lysozyme and 1 U/ml of benzonase) and incubated at 25°C with shaking at 1200 rpm for 1 hour. The cell lysates were clarified by centrifugation at 4,000 × g for 20 min at 4°C. Clarified lysates were diluted (1000-fold for β-lactamase activity 2-fold for phosphonate hydrolase activity) in order to obtain linear initial rates and measured against a single substrate concentration (90 μM for β-lactamase activity and 500 μM for phosphonate hydrolase activity).

### Purification of Strep-tagged proteins

All variants were cloned as described above, transformed, overexpressed in *E. coli* BL21 (DE3) cells and purified as described previously^27^.

### Enzyme kinetics

The kinetic parameters and activity levels of purified of enzyme variants were obtained as described previously^27^. Briefly, phosphonatase monoester hydrolase activity was monitored following the release of *p*-nitrophenol at 405 nm with an extinction coefficient of 18,300 M^−1^ cm^−1^. The β-lactamase activity was monitored at 405 nm for the Centa substrate, and molar product formation was calculated with the extinction coefficient of 6,300 M^−1^ cm^−1^.

### Thermostability assay

The thermal stability of variants was measured with a thermal shift assay as described previously^49^. Briefly, enzyme variants (2 μM) were mixed with 5 × SYPRO Orange dye (Invitrogen) in a 20l reaction and heated from 25 to 95°C in a 7500 Fast Real-Time PCR system (Applied Biosystems). Measurements were conducted in triplicate and unfolding was followed by measuring the change in fluorescence caused by binding of the dye (excitation, 488 nm; emission, 500–750 nm). The melting temperature (*T*_m_) is calculated from midpoint of the denaturation curve and values were averaged.

### Protein purification for crystallization

The NDM1 and VIM2 protein variants were expressed in *E. coli* BL21 (DE3) cells in TB medium (400 ml) supplemented with 1% glycerol, 50 μg/ml kanamycin and 200 μM ZnCl_2_. Cells were grown at 30 °C for 6 hours. The temperature was lowered to 22°C and the cells were incubated for a further 16 hours and harvested by centrifugation for 15 minutes at 8,500 × g (R9A rotor, Hitachi), then resuspended in buffer A (50 mM HEPES pH 7.5, 500 mM NaCl, 20 mM imidazole, and 200 μM ZnCl_2_) and lysed by sonication (OMNI sonic ruptor 400). Cellular debris was removed by centrifugation at 29,070 × g for 60 minutes (R15A rotor, Hitachi). The supernatant was loaded onto a 5 ml Ni-NTA superflow cartridge (Qiagen) followed by extensive washing with buffer A prior to elution of proteins in buffer B (50 mM HEPES pH 7.5, 500 mM NaCl, 500 mM imidazole, and 200 μM ZnCl_2_). Protein containing fractions were analyzed by SDS-PAGE (Bolt Mini Gels, Novex). The buffer B containing the proteins was exchanged to TEV reaction buffer (50 mM Tris-HCl pH 8.0, 0.5 mM EDTA, 1 mM DTT, and 150 mM NaCl) using HiPrep 26/10 desalting column (GE healthcare). 20% TEV (w/w) was added and incubated at 4°C for 4 days. The TEV reaction buffer was exchanged to buffer A before TEV protease and His-tag containing debris were removed by Ni-NTA superflow column (5 ml, Qiagen). His-tag cleaved protein was then concentrated using a10 kDa molecular weight cut-off MWCO ultrafiltration membrane (Amicon, Millipore) and loaded on HiLoad 16/600 Superdex 75 pg column (GE Healthcare). Proteins were eluted into crystallization buffers.

### Crystallization of NDM1-R10

NDM1-R10 protein in crystallization buffer (20 mM HEPES pH 7.5, 2 mM β-mercaptoethanol, 150 mM NaCl, and 100 μM ZnCl_2_) was concentrated to 15 mg/ml using a 10 kDa molecular weight cut-off ultrafiltration membrane (Amicon, Millipore) and crystallized by the hanging drop method. The hanging drops were prepared by mixing protein solution (1 μl) and well solution (2 μl). Crystals appeared after two weeks in 0.1 M MES (pH 6.75) and 1.3 M MgSO_4_ at 18°C and continued to grow. Crystals were soaked in cryoprotectant solution for 30 seconds (precipitant, and 25% glycerol), and flash cooled in liquid nitrogen. The MES bound crystal diffracted to 1.67 Å at beam line MX1 at the Australian Synchrotron. The product bound structure was obtained by soaking the crystal in precipitant solution, containing 15 mM substrate for 3 minutes to 30 minutes before soaking in cryoprotectant solution and flash cooling in liquid nitrogen. The crystals diffracted to 1.68-2 Å at a beam line MX2 (0.9537 Å) at the Australian Synchrotron.

### Crystallization of VIM2-R10

The first size exclusion peak of VIM2-R10 protein (dimeric fractions), in buffer containing 50 mM HEPES pH 7.5, 150 mM NaCl, and 200 μM ZnCl_2_, was concentrated to 2.6 mg/ml using a 10 kDa molecular weight cut-off ultrafiltration membrane (Amicon, Millipore) and crystallized by the hanging drop method. The drops were prepared by mixing a protein solution (2 μl), well solution (4 μl), 1 mM TCEP, and 2.5 mM p-nitrophenyl phenylphosphonate. Crystals appeared after two weeks in 0.1 M HEPES (pH 7.5) and 1.2 M sodium citrate at 18°C. Crystals were soaked in cryo-protectant solution for several minutes (precipitant, and 10% glycerol), and flash cooled in liquid nitrogen. The crystal diffracted to 2 Å at beam line MX1 (0.9537 Å) at the Australian Synchrotron.

### Data collection and structure determination

The crystallographic data were collected at 100 K at the Australian Synchrotron. Data were processed using XDS^50^. Scaling was performed using Aimless in the CCP4 program suite. Resolution estimation and data truncation were performed by using overall half-dataset correlation CC(1/2) > 0.5^51^. Molecular replacement was used to solve all structures with MOLREP^52^ using the structures deposited under PDB accession codes 3SPU and 1KO3 as starting models for NDM1 and VIM2, respectively. The model was refined using phenix.refine^53^ and Refmac v5.7^54^ in CCP4 v6.3 program^55^, and the model was subsequently optimized by iterative model building with the program COOT v0.7^56^.

### Molecular dynamics simulation

Molecular dynamics simulations VIM2 and NDM1 variants were performed using the Q simulation package^57^ and the OPLS-AA force field^58^. OPLS-AA compatible parameters for *p*-nitrophenyl phenylphosphonate (PMH substrate, PPP) were generated using MacroModel version 10.3 (Schrödinger LLC, v. 2014-1). Partial charges for PPP were calculated using the standard RESP procedure^59^, with the use of Antechamber (AmberTools 12)^60^ and Gaussian09 (Revision C.01)^61^. All other PPP parameters were as presented in the supporting information of Ref. ^62^. The structures of VIM2-WT (PDB ID 4PVO) and NDM1-WT (PDB ID 4HL2) were obtained from the Protein Data Bank, and the structure of NDM1-R10 with the PMH product bound (PDB ID 5K4M) was obtained as described in the previous section. The structures of single W93G mutants of both VIM2 and NDM1 were generated by manually mutating the respective tryptophan residues to glycine in the WT structures. In the simulations of VIM2 chain B of the PDB structure was used. The PMH substrate was placed manually in the active site of the respective enzymes based on the position of the PMH product found in the crystal structure of NDM1-R10. The Zn^2+^ ions were described using a tetrahedral dummy model based on the dummy model originally described by Åqvist and Warshel^63^. The model was built by placing four dummy atoms in a tetrahedral geometry around a central metal particle, and parametrised to reproduce the experimental solvation free energy and solvation geometry of the zinc ion (for the description of analogous octahedral dummy model and parameterization procedure see Ref.^64^). The resulting Zn^2+^ parameters used in this work are presented in **Supplementary Table 9**. All simulations were performed using surface-constrained all-atom solvent (SCAAS) boundary conditions^65^ applied to a spherical droplet of TIP3P water molecules^66^ with a radius of 24 Å centered on the bridging hydroxide ion. All protein atoms and water molecules within 85% of the solvent sphere were allowed to move freely with no restraints, atoms in the last 15% of the sphere were subject to 10 kcal mol^−1^ Å^−2^ positional restraints, and all atoms outside this sphere were subjected to 200 kcal mol^−1^ Å^−2^ positional restraints to maintain them at their crystallographic positions. Protonation states of all ionisable residues within the inner 85% of the simulation sphere were assigned using PROPKA 3.1^67,68^ and the protonation states of histidine side chains were determined by visual inspection of the surrounding hydrogen bonding pattern of each residue. The relevant protonation states and histidine protonation parameters are presented in **Supplementary Table 10**. All ionizable residues outside of the 85% of the sphere were kept in their uncharged forms to avoid simulation artefacts caused by having charged residues in the excluded region of the simulations. All systems were initially equilibrated with 200 kcal mol^−1^ Å^−2^ positional restraints over the total timescale of 95 ps, during which the alternating heating, cooling and reheating was performed to release steric clashes and equilibrate the positions of the solvent molecules and hydrogen atoms, and to reach the target simulation temperature of 300K. The initial equilibration was completed by performing 10 ns of simulation at 300K, which was followed by 100 ns production simulation, the last 50 ns of which was subject to further analysis. This protocol was repeated five times generating five independent 100ns trajectories for each system. The root mean square deviations of all C-α atoms in our simulations is shown in **Supplementary Figs. 6** and **8**, which demonstrates that all simulations have equilibrated during the 100 ns simulation time. During the final stage of the equilibration and production simulations weak 0.5 kcal mol^−1^ Å^−2^ restraints were applied on the PMH substrate in order to keep it within the simulation sphere, and 1.0 kcal mol^−1^ Å^−2^ restraints were applied on the metal ions, side chains of the metal ligands and the bridging hydroxide ion to assure proper coordination geometry of the metal centers. Apart from the very initial stages of the equilibration where a 0.01 and 0.1 fs time step was used, a 1 fs time step was used throughout the simulations. The results of the simulations were analyzed using: VMD^69^, and GROMACS^70^.

---

**Supplementary Table 1.** Information about the enzymes characterized in this study.

| Enzyme name | Uniprot ID | Organismal source | PDB ID code | Structural resolution |
| --- | --- | --- | --- | --- |
| FIM1 | K7SA42 | *Pseudomonas aeruginosa* | n.a. |  |
| EBL1 | Q2N9N3 | *Erythrobacter litoralis* | n.a. |  |
| NDM1 | C7C442 | *Klebsiella pneumonia* | 3spu | 2.1 |
| VIM2 | Q9K2N0 | *Pseudomonas aeruginosa* | 1ko3 | 1.9 |
| VIM1 | Q9XAY4 | *Pseudomonas aeruginosa* | n.a. |  |
| VIM7 | Q840P9 | *Pseudomonas aeruginosa* | 2y87 | 1.9 |

n.a, not available

**Supplementary Table 2.**
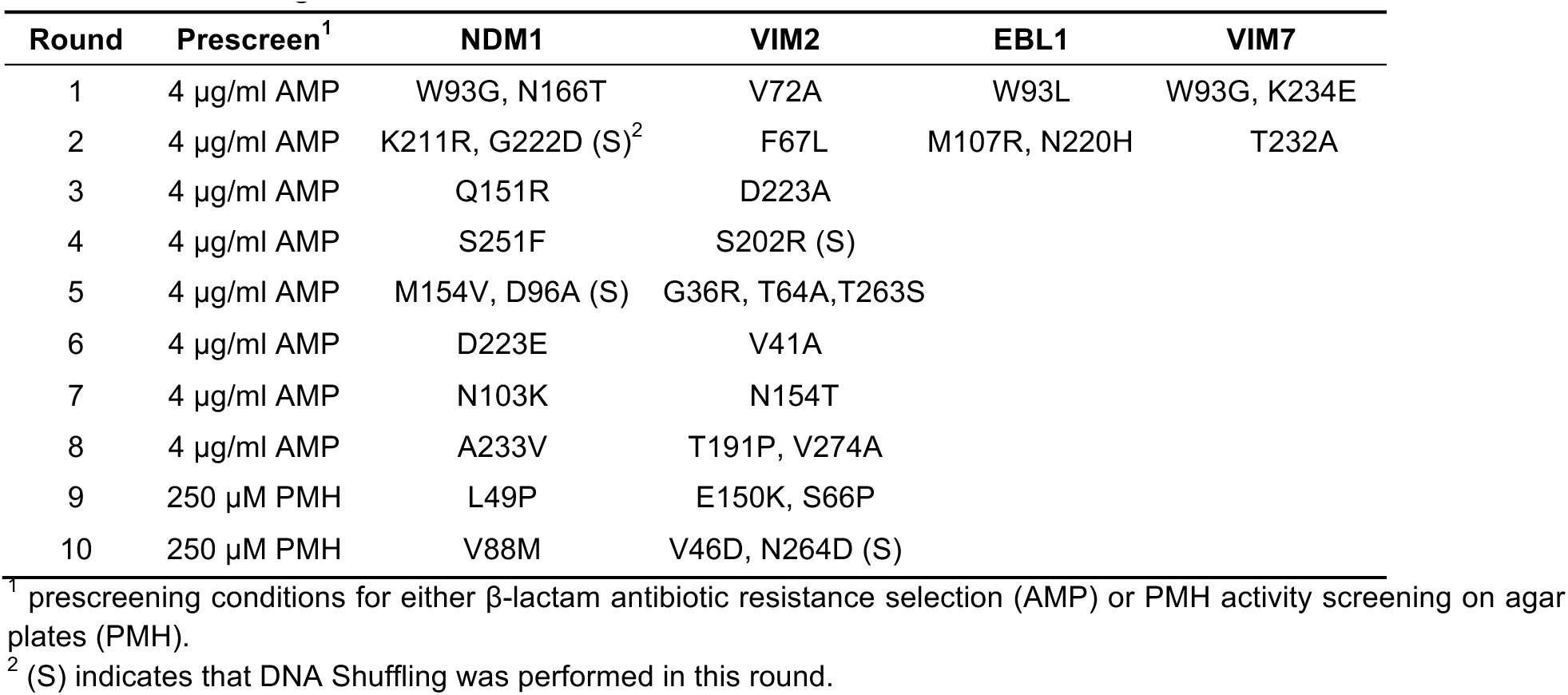
Information on the directed evolution procedure and mutations that were accumulated during directed evolution.

**Supplementary Table 3.** PMH fitness values (cell lysate activity) of evolved variants and generated mutants.

| Variant | NDM1 | VIM2 | EBL1 | VIM7 |
| --- | --- | --- | --- | --- |
|  | PMH fitness (nM/s) |  |  |  |
| WT | **0.9** ± 0.2 | **3.5** ± 0.1 | **0.1** ± 0.02 | **0.3** ± 0.02 |
| R1 | **34** ± 3 | **7.9** ± 0.3 | **4.3** ± 0.3 | **0.9** ± 0.06 |
| R2 | **190** ± 10 | **13** ± 1 | **31** ± 1 | **1.1** ± 0.1 |
| R3 | **410** ± 20 | **21** ± 1 |  |  |
| R4 | **740** ± 120 | **28** ± 1 |  |  |
| R5 | **1300** ± 150 | **50** ± 1 |  |  |
| R6 | **470** ± 60 | **70** ± 2 |  |  |
| R7 | **910** ± 30 | **70** ± 2 |  |  |
| R8 | **1800** ± 250 | **88** ± 1 |  |  |
| R9 | **1500** ± 300 | **75** ± 1 |  |  |
| R10 | **3300** ± 220 | **120** ± 1 |  |  |
| K211R/G222D | **1.2** ± 0.1 |  |  |  |
| Q151R | **1.1** ± 0.1 |  |  |  |
| S251F | **1.5** ± 0.2 |  |  |  |
| W93G | **20.7** ± 2 | 1.1 ± 0.03 |  |  |
| W93A |  | 1.3 ± 0.07 |  |  |
| W93V |  | 0.72 ± 0.06 |  |  |
| W93L |  | 0.45 ± 0.08 |  |  |
| W93F |  | 1.2 ± 0.09 |  |  |
| F67L |  | **6.5** ± 0.1 |  |  |
| D223A |  | **4.9** ± 0.3 |  |  |
| S202R |  | **6.8** ± 0.3 |  |  |

± indicates standard deviation from three independent experiments within in a 96-well plate
The cell lysate was diluted by 2-fold, and the activity was measured with 500 µM for p-nitrophenyl-phenylphosphonate. The presented value indicates the activity level in the undiluted cell lysate.

**Supplementary Table 4.** Changes in catalytic activity of purified enzymes compared to its wild-type enzyme, and melting temperature of MBL mutants.

| Enzyme | Variant | Fitness change<br>relative to WT | | $T_{m50}$ (°C) |
| --- | --- | --- | --- | --- |
| | | PMH | β-lactamase | |
| **FIM1** | WT | **1.0** | **1.0** | **53.5 ± 0.7** |
| **FIM1** | H72V | **0.8** | **3.6** | **54.0 ± 0.9** |
| **FIM1** | H72A | n.d. | n.d. | n.d. |
| **FIM1** | W93G | **6.5** | **0.9** | **46.2 ± 1.3** |
| **FIM1** | M67L | **0.8** | **0.6** | **55.0 ± 0.6** |
| **FIM1** | M67F | **1.3** | **41** | **54.2 ± 0.7** |
| **EBL1** | WT | **1.0** | **1.0** | **66.2 ± 0.5** |
| **EBL1** | W93G | **5.0** | **2.3** | **64.0 ± 0.8** |
| **EBL1** | P72V | **0.4** | **1.6** | **64.8 ± 0.6** |
| **EBL1** | L67F | **0.3** | **15.2** | **65.3 ± 1.4** |
| **EBL1** | P72A | **0.5** | **1.3** | **65.9 ± 0.6** |
| **NDM1** | WT | **1.0** | **1.0** | **55.0 ± 0.4** |
| **NDM1** | W93G | **95** | **2.0** | **47.4 ± 0.4** |
| **NDM1** | M67F | **0.5** | **1.0** | **53.9 ± 0.7** |
| **NDM1** | A72V | **0.8** | **1.3** | **54.8 ± 0.5** |
| **NDM1** | M67L | **0.5** | **1.0** | **54.7 ± 0.8** |
| **VIM2** | WT | **1.0** | **1.0** | **58.7 ± 0.6** |
| **VIM2** | V72A | **3.0** | **1.1** | **57.3 ± 0.6** |
| **VIM2** | W93G | **0.1** | **0.1** | **62.0 ± 1.0** |
| **VIM2** | F67L | **2.3** | **0.4** | **60.3 ± 0.6** |
| **VIM1** | WT | **1.0** | **1.0** | **59.0 ± 0.3** |
| **VIM1** | F67L | **1.2** | **0.6** | **58.1 ± 0.6** |
| **VIM1** | V72A | **3.0** | **1.1** | **55.1 ± 0.5** |
| **VIM1** | W93G | **0.3** | **0.1** | **55.4 ± 1.8** |
| **VIM7** | WT | **1.2** | **0.4** | **56.8 ± 1.0** |
| **VIM7** | L67F | **0.7** | **0.6** | **55.5 ± 0.6** |
| **VIM7** | V72A | **1.4** | **0.3** | **54.1 ± 0.5** |
| **VIM7** | W93G | **4.0** | **0.1** | **53.2 ± 0.8** |

n.d, not determined
± indicates standard deviation from three independent experiments
All the MBL mutants were purified using Strep-tactin affinity chromatography, and activity levels of purified enzymes were measured at single enzyme (1 μM for PMH and 1 nM for β-lactamase activity) and substrate (500 μM for PMH and 100 μM for β-lactamase activity) concentration.

**Supplementary Table 5.** Catalytic parameters of VIM2 variants.

| Variant | PMH activity | | | β-lactamase activity | | |
| --- | --- | --- | --- | --- | --- | --- |
| | $k_{\text{cat}}$ [s<sup>-1</sup>] | $K_M$ [μM] | $k_{\text{cat}} K_M$ [s<sup>-1</sup>M<sup>-1</sup>] | $k_{\text{cat}}$ [s<sup>-1</sup>] | $K_M$ [μM] | $k_{\text{cat}} K_M$ [s<sup>-1</sup>M<sup>-1</sup>] |
| WT | **0.005** ± 0.0001 | **840** ± 50 | **5.9** $\times 10^0$ | **16** ± 0.7 | **4.2** ± 0.9 | **3.8** $\times 10^6$ |
| R1 | **0.03** ± 0.001 | **2100** ± 110 | **1.3** $\times 10^1$ | **17** ± 1 | **4.7** ± 1.3 | **3.7** $\times 10^6$ |
| R2 | **0.02** ± 0.001 | **1500** ± 100 | **1.6** $\times 10^1$ | **23** ± 1 | **9.6** ± 2.3 | **2.4** $\times 10^6$ |
| R3 | **0.04** ± 0.0005 | **850** ± 30 | **4.8** $\times 10^1$ | **37** ± 2 | **100** ± 21 | **3.7** $\times 10^5$ |
| R4 | **0.07** ± 0.001 | **1200** ± 100 | **5.8** $\times 10^1$ | **32** ± 3 | **21** ± 6.1 | **1.5** $\times 10^6$ |
| R5 | **0.09** ± 0.002 | **640** ± 30 | **1.4** $\times 10^2$ | **53** ± 1 | **37** ± 3.5 | **1.4** $\times 10^6$ |
| R6 | **0.15** ± 0.004 | **880** ± 70 | **1.8** $\times 10^2$ | **40** ± 6 | **110** ± 46 | **3.7** $\times 10^5$ |
| R7 | **0.08** ± 0.003 | **890** ± 90 | **9.3** $\times 10^1$ | **17** ± 2 | **32** ± 11 | **5.4** $\times 10^5$ |
| R8 | **0.15** ± 0.01 | **890** ± 80 | **1.6** $\times 10^2$ | **33** ± 4 | **55** ± 19 | **6.0** $\times 10^5$ |
| R9 | **0.15** ± 0.007 | **1300** ± 200 | **1.2** $\times 10^2$ | **19** ± 0.3 | **33** ± 3 | **5.8** $\times 10^5$ |
| R10 | **0.15** ± 0.005 | **1000** ± 100 | **1.4** $\times 10^2$ | **15** ± 0.3 | **34** ± 3 | **4.6** $\times 10^5$ |

± indicates standard deviation from three independent measurements

**Supplementary Table 6.**
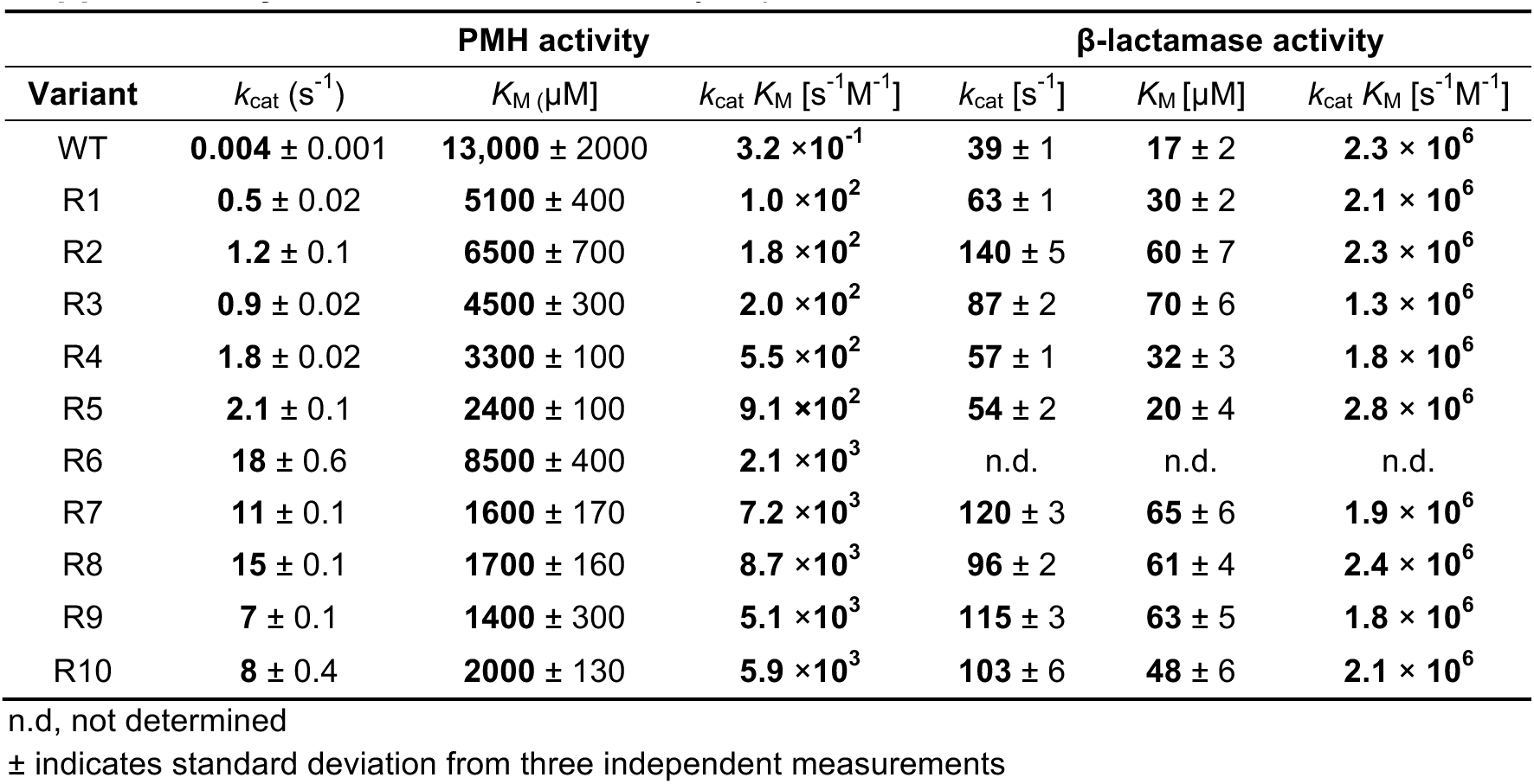
Individual catalytic parameters of NDM1 variants.

**Supplementary Table 7.** Solubility and melting temperature of NDM1 and VIM2 variants.

| **Variant** | **NDM1** |  | **VIM2** |  |
| --- | --- | --- | --- | --- |
| | Solubility (%) | $T_m$ (°C) | Solubility (%) | $T_m$ (°C) |
| WT | 43 | **55.0** ± 0.4 | 40 | **58.7** ± 0.6 |
| R1 | 25 | **48.1** ± 0.6 | 44 | **57.3** ± 0.6 |
| R2 | 21 | **51.0** ± 0.5 | 42 | **60.3** ± 2.1 |
| R3 | 27 | **52.4** ± 0.8 | 37 | **56.3** ± 1.5 |
| R4 | 16 | **53.1** ± 1.1 | 48 | **58.0** ± 0.1 |
| R5 | 18 | **51.9** ± 0.3 | 57 | **54.3** ± 2.3 |
| R6 | 18 | **50.1** ± 0.8 | 62 | **54.3** ± 2.3 |
| R7 | 21 | **51.0** ± 0.7 | 61 | **55.3** ± 1.2 |
| R8 | 28 | **55.8** ± 0.8 | 69 | **61.7** ± 0.6 |
| R9 | 16 | **54.1** ± 0.4 | 86 | **55.3** ± 0.6 |
| R10 | 34 | **55.0** ± 0.9 | 67 | **55.3** ± 0.6 |

± indicates standard deviation from three independent experiments

**Supplementary Table 8.** Crystallographic data collection and refinement statistics.

| Variant | NDM1-R10 | NDM1-R10 (product) | VIM2-R10 |
| --- | --- | --- | --- |
| PDB ID | 5JQJ | 5K4M | 6BM9 |
| Wavelength (Å) | 0.9537 | 0.9537 | 0.9537 |
| No. copies in an asymmetric unit | 1 | 1 | 4 |
| Resolution range (Å) | 34.45-1.67<br>(1.73-1.67) | 39.13-1.98<br>(2.05-1.98) | 37.89-2.19<br>(2.27-2.19) |
| Space group | C 2 2 2<sub>1</sub> | C 2 2 2<sub>1</sub> | C 1 2 1 |
| Unit cell (Å, °) | 37.80 137.72 77.46 90<br>90 90 | 37.82 138.16 78.20 90<br>90 90 | 128.60 41.67 156.76<br>90 99 90 |
| Total reflections | 46608 (3668) | 29257 (2838) | 150766 (13607) |
| Unique reflections | 23318 (1837) | 14613 (1421) | 41963 (4025) |
| Multiplicity | 2.0 (2.0) | 2.0 (2.0) | 3.6 (3.4) |
| Completeness (%) | 97.22 (77.72) | 99.72 (99.37) | 97.82 (96.08) |
| Mean I/sigma(I) | 17.93 (1.89) | 11.13 (5.78) | 11.28 (1.86) |
| Wilson B-factor (Å<sup>2</sup>) | 19.27 | 14.37 | 32.54 |
| <sup>a</sup>R-merge | 0.029 (0.46) | 0.029 (0.08) | 0.094 (0.76) |
| <sup>b</sup>R-meas | 0.04 | 0.04 | 0.11 |
| <sup>c</sup>CC1/2 | 0.999 (0.588) | 0.999 (0.978) | 0.996 (0.609) |
| <sup>d</sup>CC* | 1.00 (0.861) | 1.00 (0.994) | 0.999 (0.870) |
| R-work | 0.144 (0.240) | 0.142 (0.150) | 0.223 (0.323) |
| R-free | 0.187 (0.287) | 0.208 (0.198) | 0.268 (0.337) |
| Number of non-hydrogen atoms | 2052 | 1982 | 7053 |
| macromolecules | 1758 | 1749 | 6842 |
| ligands | 28 | 46 | 16 |
| water | 266 | 187 | 195 |
| Protein residues | 230 | 230 | 902 |
| RMS(bonds) (Å) | 0.02 | 0.017 | 0.003 |
| RMS(angles) (°) | 2.08 | 1.95 | 0.76 |
| Ramachandran preferred (%) | 98.05 | 96.63 | 95.71 |
| Ramachandran allowed (%) | 0.98 | 2.4 | 3.50 |
| Ramachandran outliers (%) | 0.98 | 0.96 | 0.79 |
| Clashscore | 3.95 | 5.67 | 4.94 |
| Average B-factor (Å<sup>2</sup>) | 24 | 16.8 | 45.1 |
| macromolecules | 22.1 | 15.4 | 45.2 |
| ligands | 36.8 | 29.7 | 57.3 |
| solvent | 35 | 26.6 | 40.9 |

\* Highest-resolution shell is shown in parentheses.

**Supplementary Table 9.** Parameters used to describe the Zn^2+^ ions in our MD simulations.

| bond type<sup>a</sup> | | $K_b$ | $r_0$ | |
| --- | --- | --- | --- | --- |
| Zn-D |  | 800.0 | 0.90 |  |
| D-D |  | 800.0 | 1.47 |  |
| angle type<sup>b</sup> | | $K_\theta$ | $\theta_0$ | |
| D-Zn-D |  | 250.0 | 109.50 |  |
| D-D-Zn |  | 250.0 | 32.25 |  |
| D-D-D |  | 250.0 | 60.00 |  |
| atom type | mass | charge | $A_i^c$ | $B_i^c$ |
| Zn | 53.4 | 0.00<sup>d</sup>/-1.00<sup>e</sup> | 41.00 | 32.00 |
| D | 3.0 | 0.50<sup>d</sup>/0.75<sup>e</sup> | 0.05 | 0.05<sup>d</sup>/0.00<sup>e</sup> |

**Supplementary Table 10.** List of relevant ionized states as well as the protonation patterns of histidine residues in our molecular dynamics simulations. All other residues were kept in their unionized forms as they were outside the simulation sphere (see main text).

| Residue | Residue number |  |  |
| --- | --- | --- | --- |
|  | NDM1 WT and W93G | NDM1 R10 | VIM2 |
| Asp | 66, 90, 95, 124, 130, 192, 212, 223, 225, 254 | 66, 90, 95, 124, 130, 192, 212, 222, 225, 254 | 62, 84, 119, 120, 126, 199, 236, 238 |
| Glu | 152 | 152, 223 | 147, 149, 166, 225 |
| Lys | 125, 211, 216 | 125, 216 | 90 |
| Arg | - | 151, 211 | 121, 129, 143, 144, 228, 296 |
| His-δ | 250, 133, 120, 189 | 250, 133, 120, 189 | 116, 196, 263 |
| His-ε | 61, 122, 159 | 61, 122, 159 | 55, 118, 170, 252, 285, 293 |
| Hip | 228, 261 | 228, 261 | - |

## References

1. Dworkin, I. Uncovering cryptic genetic variation. Nat Rev Genet 5, 681–690 (2004).

2. Paaby, A. B. & Rockman, M. V. Cryptic genetic variation: evolution’s hidden substrate. Nat Rev Genet 15, 247–258 (2014).

3. Le Rouzic, A., Lerouzic, A., Carlborg, O. & Carlborg, O. Evolutionary potential of hidden genetic variation. Trends in Ecology & Evolution 23, 33–37 (2008).

4. Specchia, V. et al. Hsp90 prevents phenotypic variation by suppressing the mutagenic activity of transposons. Nature 463, 662–665 (2010).

5. Hayden, E. J., Ferrada, E. & Wagner, A. Cryptic genetic variation promotes rapid evolutionary adaptation in an RNA enzyme. Nature 474, 92–95 (2011).

6. Amitai, G., Gupta, R. D. & Tawfik, D. S. Latent evolutionary potentials under the neutral mutational drift of an enzyme. HFSP J. 1, 67–78 (2007).

7. Bloom, J. D., Romero, P. A., Lu, Z. & Arnold, F. H. Neutral genetic drift can alter promiscuous protein functions, potentially aiding functional evolution. Biol Direct 2, 17 (2007).

8. Rohner, N. et al. Cryptic variation in morphological evolution: HSP90 as a capacitor for loss of eyes in cavefish. Science 342, 1372–1375 (2013).

9. Wagner, A. Neutralism and selectionism: a network-based reconciliation. Nat Rev Genet 9, 965–974 (2008).

10. Romero, P. A. & Arnold, F. H. Exploring protein fitness landscapes by directed evolution. Nat Rev Mol Cell Biol 10, 866–876 (2009).

11. de Visser, J. A. G. M. & Krug, J. Empirical fitness landscapes and the predictability of evolution. Nat Rev Genet 15, 480–490 (2014).

12. Starr, T. N. & Thornton, J. W. Epistasis in protein evolution. Protein Sci 25, 1204–1218 (2016).

13. Miton, C. M. & Tokuriki, N. How mutational epistasis impairs predictability in protein evolution and design. Protein Sci 25, 1260–1272 (2016).

14. Parera, M. & Martínez, M. A. Strong epistatic interactions within a single protein. Molecular Biology and Evolution 31, 1546–1553 (2014).

15. Kershner, J. P. & Copley, S. D. Differential effects of a mutation on the normal and promiscuous activities of orthologs: implications for natural and directed evolution. 32, 100–108 (2015).

16. Harms, M. J., Thornton, J. W. & Thornton, J. W. Historical contingency and its biophysical basis in glucocorticoid receptor evolution. Nature 512, 203–207 (2014).

17. Starr, T. N., Picton, L. K. & Thornton, J. W. Alternative evolutionary histories in the sequence space of an ancient protein. Nature 549, 409–413 (2017).

18. O’Loughlin, T. L. Natural history as a predictor of protein evolvability. Protein Eng Des Sel 19, 439–442 (2006).

19. Burch, C. L. & Chao, L. Evolvability of an RNA virus is determined by its mutational neighbourhood. Nature 406, 625–628 (2000).

20. Yip, S. H. C. & Matsumura, I. Substrate ambiguous enzymes within the *Escherichia coli* proteome offer different evolutionary solutions to the same problem. Mol Biol Evol 30, 2001–2012 (2013).

21. Schulenburg, C., Stark, Y., Künzle, M. & Hilvert, D. Comparative laboratory evolution of ordered and disordered enzymes. J Biol Chem 290, 9310–9320 (2015).

22. Toth-Petroczy, A. & Tawfik, D. S. The robustness and innovability of protein folds. Curr Opin Struct Biol 26, 131–138 (2014).

23. Bloom, J. D. & Arnold, F. H. In the light of directed evolution: pathways of adaptive protein evolution. Proc Natl Acad Sci U S A 106 Suppl 1, 9995–10000 (2009).

24. Tokuriki, N. & Tawfik, D. S. Stability effects of mutations and protein evolvability. Curr Opin StructBiol 19, 596–604 (2009).

25. De Pristo, M. A., Weinreich, D. M. & Hartl, D. L. Missense meanderings in sequence space: a biophysical view of protein evolution. Nat Rev Genet 6, 678–687 (2005).

26. Bebrone, C. Metallo-β-lactamases (classification, activity, genetic organization, structure, zinc coordination) and their superfamily. Biochem Pharmacol 74, 1686–1701 (2007).

27. Baier, F. & Tokuriki, N. Connectivity between catalytic landscapes of the metallo-β-lactamase superfamily. J Mol Biol 426, 1–15 (2014).

28. Baier, F., Chen, J., Solomonson, M., Strynadka, N. C. J. & Tokuriki, N. Distinct metal isoforms underlie promiscuous activity profiles of metalloenzymes. ACS Chem Biol 10, 1684–1693 (2015).

29. Hartl, D. L. What can we learn from fitness landscapes? Curr Opin Microbiol 21, 51–57 (2014).

30. Tokuriki, N., Jackson, C. J. & Tawfik, D. S. Diminishing returns and tradeoffs constrain the laboratory optimization of an enzyme. Nature Commun 3, 1257 (2012).

31. King, D. T., Worrall, L. J., Gruninger, R. & Strynadka, N. C. J. New Delhi metallo-β-lactamase: Structural insights into β-lactam recognition and inhibition. J Am Chem Soc 134, 11362–11365 (2012).

32. Garcia-Saez, I., Docquier, J. D., Rossolini, G. M. & Dideberg, O. The threedimensional structure of VIM-2, a Zn-β-lactamase from *Pseudomonas aeruginosa* in its reduced and oxidised form. J Mol Biol 375, 604–611 (2008).

33. Aharoni, A. et al. The ‘evolvability’ of promiscuous protein functions. Nat Genet 268, 27458–76 (2004).

34. Baier, F., Copp, J. N. & Tokuriki, N. Evolution of enzyme superfamilies: comprehensive exploration of sequence-function relationships. Biochemistry 55, 6375–6388 (2016).

35. Weinreich, D. M. Darwinian evolution can follow only very few mutational paths to fitter proteins. Science 312, 111–114 (2006).

36. Salverda, M. L. M. et al. Initial mutations direct alternative pathways of protein evolution. PLoS Genet 7, e1001321 (2011).

37. Lobkovsky, A. E. & Koonin, E. V. Replaying the tape of life: quantification of the predictability of evolution. 3, 246–246 (2012).

38. Dickinson, B. C., Leconte, A. M., Allen, B., Esvelt, K. M. & Liu, D. R. Experimental interrogation of the path dependence and stochasticity of protein evolution using phage-assisted continuous evolution. Proc Natl Acad Sci U S A 110, 9007–9012 (2013).

39. Palmer, A. C., Bershtein, S. & Kishony, R. Delayed commitment to evolutionary fate in antibiotic resistance fitness landscapes. Nature Commun 6, 7385 (2015).

40. Kaltenbach, M., Jackson, C. J., Campbell, E. C., Hollfelder, F. & Tokuriki, N. Reverse evolution leads to genotypic incompatibility despite functional and active site convergence. eLife 4, (2015).

41. Thornton, J. W. Evolutionary biochemistry: revealing the historical and physical causes of protein properties. Nat Rev Genet 14, 559–571 (2013).

42. Kaltenbach, M. & Tokuriki, N. Dynamics and constraints of enzyme evolution. J. Exp. Zool. B Mol. Dev. Evol. 322, 468–487 (2014).

43. Liang, B. et al. Facilitation of bacterial adaptation to chlorothalonil-contaminated sites by horizontal transfer of the chlorothalonil hydrolytic dehalogenase gene. Appl Environ Microbiol 77, 4268–4272 (2011).

44. Singh, B. K. Organophosphorus-degrading bacteria: ecology and industrial applications. Nat Rev Microbiol 7, 156–164 (2008).

45. Dalby, P. A. Strategy and success for the directed evolution of enzymes. Curr Opin in Struct Biol 21, 473–480 (2011).

46. Davids, T., Schmidt, M., Böttcher, D. & Bornscheuer, U. T. Strategies for the discovery and engineering of enzymes for biocatalysis. Curr Opin Chem Biol 17, 215–220 (2013).

47. Goldsmith, M., Ashani, Y., Leader, H. & Sussman, J. L. Overcoming an optimization plateau in the directed evolution of highly efficient nerve agent bioscavengers. Protein Eng Des Sel 30, 333–345 (2017).

## Supplementary References

48. Zhao, H., Giver, L., Shao, Z., Affholter, J. A. & Arnold, F. H. Molecular evolution by staggered extension process (StEP) in vitro recombination. Nat Biotechnol 16, 258–261 (1998).

49. Kaltenbach, M. & Tokuriki, N. GroEL/ES buffering and compensatory mutations promote protein evolution by stabilizing folding intermediates. J Mol Biol 425, 3403–3414 (2013).

50. Kabsch, W. XDS. Acta Crystallogr D Biol Crystallogr 66, 125–132 (2010).

51. Karplus, P. A. & Diederichs, K. Linking crystallographic model and data quality. Science 336, 1030–1033 (2012).

52. Vagin, A. & Teplyakov, A. Molecular replacement with MOLREP. Acta Crystallogr D Biol Crystallogr 66, 22–25 (2010).

53. Afonine, P. V. et al. Towards automated crystallographic structure refinement with phenix.refine. Acta Crystallogr D Biol Crystallogr 68, 352–367 (2012).

54. Murshudov, G. N. et al. REFMAC5 for the refinement of macromolecular crystal structures. Acta Crystallogr D Biol Crystallogr 67, 355–367 (2011).

55. Winn, M. D. et al. Overview of the CCP4 suite and current developments. Acta Crystallogr D Biol Crystallogr 67, 235–242 (2011).

56. Emsley, P. & Cowtan, K. Coot: model-building tools for molecular graphics. Acta Crystallogr D Biol Crystallogr 60, 2126–2132 (2004).

57. Marelius, J. Q: a molecular dynamics program for free energy calculations and empirical valence bond simulations in biomolecular systems. J Molecular Graphics and Modelling 16, 213–225 (1998).

58. Jorgensen, W. L., Maxwell, D. S. & Tirado-Rives, J. Development and testing of the OPLS all-atom force field on conformational rnergetics and properties of organic liquids. J Am Chem Soc 118, 11225–11236 (1996).

59. Cieplak, P., Cornell, W. D., Bayly, C. & Kollman, P. A. Application of the multimolecule and multiconformational RESP methodology to biopolymers: Charge derivation for DNA, RNA, and proteins. J Comput Chem 16, 1357–1377 (1995).

60. Wang, J., Wang, W., Kollman, P. A. & Case, D. A. Automatic atom type and bond type perception in molecular mechanical calculations. J Mol Graph Model 25, 247–260 (2006).

61. Frisch, M. J. et al. Gaussian 09 C. 01. (2009).

62. Barrozo, A., Duarte, F., Bauer, P., Carvalho, A. T. P. & Kamerlin, S. C. L. Cooperative electrostatic interactions drive functional evolution in the alkaline phosphatase superfamily. J Am Chem Soc 137, 9061–9076 (2015).

63. Aqvist, J. & Warshel, A. Calculations of free energy profiles for the *Staphylococcal* nuclease catalyzed reaction. Biochemistry 28, 4680–4689 (1989).

64. Duarte, F. et al. Force field independent metal parameters using a nonbonded dummy model. J Phys Chem B 118, 4351–4362 (2014).

65. King, G. & Warshel, A. A surface constrained all-atom solvent model for effective simulations of polar solutions. J Chem Phys 91, 3647–3661 (1989).

66. Jorgensen, W. L., Chandrasekhar, J., Madura, J. D., Impey, R. W. & Klein, M. L. Comparison of simple potential functions for simulating liquid water. J Chem Phys 79, 926–935 (1983).

67. Søndergaard, C. R., Olsson, M. H. M., Rostkowski, M. & Jensen, J. H. Improved treatment of ligands and coupling effects in empirical calculation and rationalization of pKa values. J. Chem. Theory Comput. 7, 2284–2295 (2011).

68. Olsson, M. H. M., Søndergaard, C. R., Rostkowski, M. & Jensen, J. H. PROPKA3: Consistent treatment of internal and surface residues in empirical pKa predictions. J. Chem. Theory Comput. 7, 525–537 (2011).

69. Humphrey, W., Dalke, A. & Schulten, K. VMD: Visual molecular dynamics. J Mol Graph 14, 33–38 (1996).

70. Abraham, M. J. et al. GROMACS: High performance molecular simulations through multi-level parallelism from laptops to supercomputers. SoftwareX 1–2, 19–25 (2015).

71. Weiss, M. S. & Hilgenfeld, R. On the use of the merging Rfactor as a quality indicator for X-ray data. J Appl Crystallogr 30, 203–205 (1997).

72. Diederichs, K. & Karplus, P. A. Improved R-factors for diffraction data analysis in macromolecular crystallography. Nat Struct Biol 4, 269–275 (1997).

73. Diederichs, K. & Karplus, P. A. Better models by discarding data? Acta Crystallogr D Biol Crystallogr 69, 1215–1222 (2013).

